# Extracellular domains of E-cadherin determine key mechanical phenotypes of an epithelium through cell- and non-cell-autonomous outside-in signalling

**DOI:** 10.1101/2020.03.17.996181

**Authors:** D.M.K. Aladin, Y.S. Chu, R.C. Robinson, S. Dufour, V. Viasnoff, N. Borghi, J.P. Thiery

## Abstract

Cadherins control intercellular adhesion in most metazoans. In vertebrates, intercellular adhesion differs considerably between cadherins of type-I and type-II, predominantly due to their different extracellular regions. Yet, intercellular adhesion critically depends on actomyosin contractility, in which the role of the cadherin extracellular region is unclear. Here, we dissect the roles of the Extracellular Cadherin (EC) Ig-like domains by expressing chimeric E-cadherin with E-cadherin and cadherin-7 Ig-like domains in cells naturally devoid of cadherins. Using cell-cell separation, cortical tension measurement, tissue-scale stretching and migration assays, we show that distinct EC repeats in the extracellular region of cadherins differentially modulate epithelial sheet integrity, cell-cell separation forces, and cell cortical tension through a Cdc42 pathway, which further differentially regulate epithelial tensile strength, ductility, and ultimately collective migration. Interestingly, dissipative processes rather than static adhesion energy mostly dominate cell-cell separation forces. We provide a framework for the emergence of epithelial phenotypes from cell mechanical properties dependent on EC outside-in signalling.

---

Intercellular adhesion plays an important role in the development and maintenance of multicellular organisms^1-3^. Such adhesions are established and maintained through calcium-dependent, cell-cell adhesion molecules known as cadherins. Type-I and type-II cadherins, also known as classical cadherins, form one of six highly conserved branches of the cadherin superfamily, classified based on protein sequences in the extracellular cadherin (EC) domains. Classical cadherins typically have an extracellular region comprising five EC repeats (EC1-5), a single transmembrane region, and a cytoplasmic region^4,5^, and are often found concentrated in adherens junctions, dynamically interacting with the contractile cytoskeleton through catenin complexes. Despite striking structural similarities, E-cadherin (type-I) engenders robust intercellular adhesion, as seen in the epithelium, whereas cadherin-7 (type-II) is associated with much weaker adhesions, such as that in the mesenchyme^6,7^. This phenotypic difference may be attributed to the cadherin extracellular region. Indeed, whereas all three regions of cadherins are essential for normal function, the EC region dictates differences in cell-cell separation forces (SFs)—a metric that may be used as a proxy for intercellular adhesion energy^8^—between type-I and type-II cadherin-expressing cell pairs^6^.

In E-cadherin (E-cad) EC domains, henceforth referred to as EECs, the first and outermost EC domain (EEC1) has a tryptophan residue at position 2 (Trp2), which interacts with a hydrophobic pocket in the opposing EEC1 of its cadherin pair to form strand-swap dimers. In contrast, in cadherin-7 (cad-7) EC domains, henceforth referred to as 7ECs, the conserved Trp2 and Trp4 residues in 7EC1 anchor into a larger hydrophobic pocket to form two strand-swaps^9^. Associations and dissociations of strand-swapped dimers were recently identified to be mediated by binding intermediates referred to as X-dimers, which formed through extensive surface interactions between the residues near the EC1-2 calcium binding sites^10-12^. X-dimers behave as catch bonds, where the bond lifetime increases in the presence of tensile forces^13^; comparatively, strand-swap dimers form slip bonds, for which the bond lifetime decreases under tensile force. Structural and biophysical studies corroborate that cadherins tune their *trans*-bond lifetimes by switching between X- and strand-swap dimer states^14^. As both type-I and type-II cadherins are known to form X-dimers, whether the strand-swapping specifics of type-I and –II cadherins, or other causes are responsible for the significant difference between E-cad and cad-7 mediated cell-cell SFs remains to be uncovered^10,15^.

Indeed, although the residues involved in strand-swap and X dimerizations are exclusive to EC1,2 domains, mounting evidence suggests that the modular cadherin structure can also engage in multiple, *trans*- and *cis-*bonds involving more than the outer two EC Ig-like domains^16-24^. Moreover, *in silico* studies show that EEC and 7EC domains unfold in remarkably different ways under mechanical forces^25^. This raises the possibility that differences in adhesive properties between type-I and type-II cadherins arise from other ECs than EC1,2, and do not rely uniquely on strand-swap or X dimerizations.

Dual pipette cell-cell SF assays revealed that, after a few minutes of initial cell-cell contact, cell-cell SF is crucially dependent on the actin cytoskeletal anchorage of cadherins^26^, in a cadherin type-dependent manner^6^, and this is further supported by observations that cadherins uncouple from the cytoskeleton rather than from each other in zebrafish embryo cell-cell separation^27^. Indeed, cadherins appear to contribute to intercellular adhesion energy by locally down-regulating actomyosin cortical contractility^28-30^, with marginal contributions from the molecular binding energy of *trans*-interacting E-cad between cells^27,31,32^. Nevertheless, myosin-II–driven actin dynamics do modulate the immobilization of E-cad at the cell-cell contact periphery in a mechanosensitive manner^33^, where cadherin *trans*-interactions are required to physically link cells together^27,29^. Whether specific ECs of type-I and -II cadherins differentially regulate the cortical actomyosin cytoskeleton, and whether such a regulation depends on trans-interactions remain to be demonstrated.

In this work, we thus sought to address how the distinct ECs of type-I and -II cadherins regulate cell-cell adhesive properties, the contributions of trans-interactions and their effects on cortical mechanics, and the consequences at the tissue level. To do so, we examined the properties of cells individually expressing four different chimeric cadherins whose five EC regions are combinations of EC domains from E-cad or cad-7. Each chimera retained the E-cad transmembrane and cytoplasmic regions to preserve the interactions of E-cad, the catenin complex, and the cytoskeleton. We found that epithelial-like phenotype remained after exchanging EEC1-3 to 7EC1-3 but was fully lost upon exchange of the 4 first domains. Using a novel pillar-pipette assay to measure cell-cell SFs, we found that the exchange of the sole EEC3 by 7-EC3 lead to a very strong decrease of SF, and the replacement of the 3 domains EEC1-3 drastically reduced SF maturation. Next, we determined by pipette aspiration that EEC4 and EEC1-3 antagonistically affected CdC42 activity and cell cortical tension cell-autonomously. From this, and the measurement of cell-cell compaction, we determined that the static adhesion energy was a minor contributor to cell-cell SF, which was therefore mostly governed by dissipative processes to which EC region stretching before trans-bond rupture quantitatively suffices to contribute. Finally, using a custom substrate-free tissue stretcher, we provide evidence that EEC3 offers tensile strength to cell sheets, while EEC1-3 provides ductility, in accordance with their respective effects on cell-cell SF and cortical tension. We propose that the antagonistic effects of tissue ductility and cohesion endowed by the different EC regions ultimately determines the speed of collective cell migration.

## RESULTS

### Cadherin/catenin complex recruitment and epithelial integrity is largely independent of EEC1-3 (but not of EEC1-4) domains

To study the activities of the EEC domains, we designed chimeric mouse E-cad–cad-7 ectodomains — (N)77EEE(C), 777EE, EE7EE and 7777E—connected to the transmembrane and cytoplasmic regions of E-cad tagged with eGFP at the C-terminus (Fig. 1a,b). Computer models of juxtaposed domains at the hybrid interfaces showed similar domain orientations between chimeras at each interface (e.g. compare 77EEE versus EE777) suggesting that the design of the chimeras did not disrupt domain:domain interactions. Furthermore, analysis of the calcium-binding site demonstrated that at least three sidechains were available to coordinate each calcium ion, supporting that the domain interfaces are stabilized by calcium in a similar manner to the native proteins (Fig. 1c)^34^. These chimeras and wild-type E-cad (EEEEE) were stably expressed individually in S180 cells using a lentivirus-based transduction technique, selected by Puromycin resistance and sorted using fluorescence activated cell-sorting (FACS) to pool eGFP positive cells (Supplementary Fig. 1). S180 untransfected cells were used as a control.

**Figure 1:**
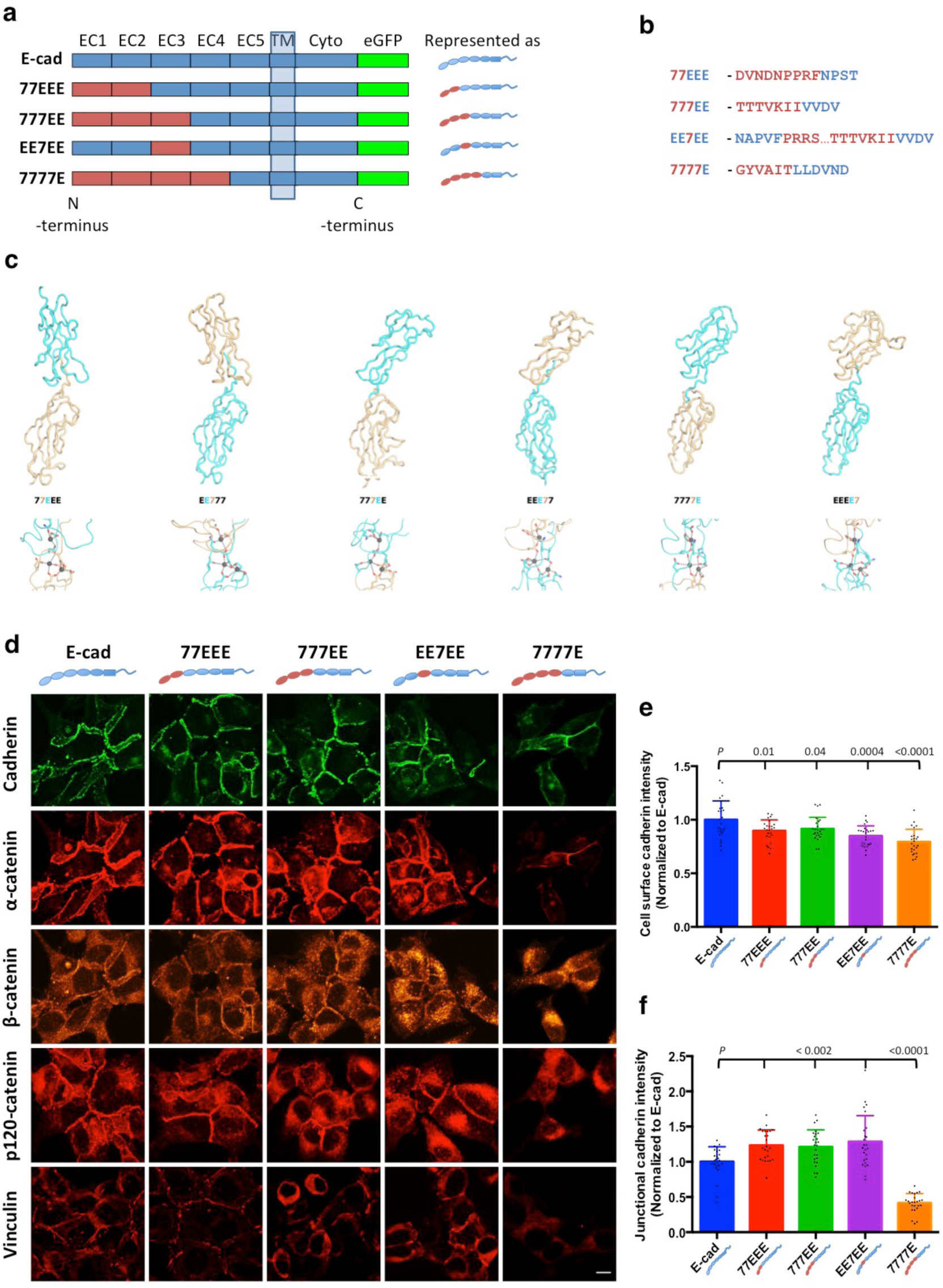
Wild-type and chimeric cadherins used in this study and their colocalization with catenin complexes. **(a)** Schematic representation of the wild-type E-cadherin (E-cad) and chimeras with the EEC1-EEC2, EEC1-EEC3, EEC3 and EEC1-EEC4 swapped with the respective 7EC domains. The EC5 domain, transmembrane domain (TM) and cytoplasmic (Cyto) domain of all chimeras are of E-cad. Wild-type E-cad and chimeras are tagged with eGFP at the C-terminus. The cartoon on the right shows the arrangement of E-cad and cad-7 domains in the wild-type and chimeric cadherins; this will be used in all subsequent figures for easy reference. **(b)** Amino acid sequence at the E-cad–cad-7 chimeric interface. Note-calcium binding sites are preserved. **(c)** Computer models of juxtaposed domains at the hybrid interfaces. Upper panel shows models of the hybrid interfaces between cadherin--7 (7, gold) and E--cadherin (E, cyan). Lower panel shows residues coordinating the predicted calcium ions. Main chain interactions with the calcium ions are not included for clarity. **(d)** Immunofluorescence microscopy of S180 cells expressing eGFP-tagged wild-type E-cad, 77EEE, 777EE, EE7EE and 7777E chimeras, stained with mouse anti-α-catenin (α-catenin), anti-β-catenin (β-catenin), anti-p120-catenin (p120-catenin) and anti-vinculin (Vinculin) antibodies (presented in their original intensities). Note the colocalization of cadherins, α-, β-, and p120-catenins and vinculin at the cell-cell junctions in E-cad, 77EEE, 777EE and EE7EE expressing cells. In the 7777E-expressing cells, cadherin and α-catenin intensity are low at junctions when compared with other cells. β- and p120-catenins in 7777E-expressing cells are more diffusely distributed along the junctions. Vinculin colocalization is not observed in 7777E-expressing cells. Cytoplasmic vinculin is more abundant in 777EE- and EE7EE-expressing cells than in E-cad- and 77EEE-expressing cells. Scale bar, 10 µm. Note: images of cadherin, α-catenin and β-catenin are acquired from same samples. **(e)** Cadherin intensity at the cell surface quantified from confocal images acquired at the cell-substrate interface. Note that the level of E-cad at cell surface is significantly higher than that of the chimeric cadherins (*n* = 25 in each group, *n* is the number of cells analyzed). **(f)** Histogram of junctional cadherin intensity normalized to junctional E-cad intensity. Note that the intensity of 77EEE, 777EE and EE7EE are significantly higher than that of E-cad; the intensity of 7777E is significantly lower than E-cad (*n* = 25 in each group, *n* is the number of junctions analyzed).

We examined the colocalization of junctional cadherins with catenins and vinculin by immunofluorescence labelling (Fig. 1d). We observed that α-, β- and p120-catenins were expressed in all clones and co-localized with cadherins in E-cad, 77EEE-, 777EE- and EE7EE-expressing clones. In 7777E-expressing cells, however, catenins were more diffusely distributed (Fig. 1d). Vinculin expression showed a similar trend (Fig. 1d).

Although the cells were sorted by FACS based on overall eGFP signal, cadherin surface levels are an important determinant of the maximum cell-cell SF and its maturation kinetics^6^. As we have used chimeric cadherin ectodomains in this study, the option of using antibodies targeting the ectodomains for labeling cell surface cadherins for FACS sorting did not seem feasible. Therefore, to determine the cell surface cadherin levels, we quantified eGFP intensity at the cell surfaces in contact with the substrate using confocal microscopy on fixed samples (Supplementary Fig. 2a-e). The intensities of cell-surface chimeric cadherins were normalized to E-cad intensity. The surface levels of all chimeric cadherins were significantly lower (*P* ≤ 0.04), but no more than 20%, than those of E-cad (Fig. 1e and Supplementary Fig. 2a-e). We also determined cadherin-eGFP intensity at cell-cell junctions in 2D culture by confocal microscopy on fixed samples, and found that junctional cadherin-eGFP intensities in 77EEE, 777EE and EE7EE clones were significantly higher than that of E-cad clones (*P* < 0.002; Fig. 1f). However, junctional levels of 7777E was significantly lower than that of all the other groups (*P* < 0.0001; Fig. 1f). Furthermore, the difference between the junctional levels of 7777E and E-cad was much larger than that between surface levels of the same group pair, or that between junctional levels of E-cad and any other chimeric cadherin. Together, these results show that the cadherin, catenin, and vinculin junctional recruitment hardly depends on EEC1-3. It suggests that the specifics of strand-swapping of type-I or -II cadherins may not be involved. To verify the importance of strand-swapping, we stably expressed strand-swap incompetent, full-length E-cad mutants (W2A) tagged with eGFP in S180 cells using the lentivirus-based transduction technique. These cells did not form stable junctions in 2D cultures (Supplementary Movie 1, left panel), similar to the 7777E cells. Thus, epithelial integrity required strand-swapping *per se* independently of the cadherin-type. We thus sought to determine the specific roles of E-cad EC1-3.

### EEC3 accounts for half the strength of mature cell-cell separation force and synergizes with EEC1,2, for fast junction maturation

To quantify the SF of cell-cell junctions, we used a novel pillar-pipette assay, where one of the pipettes of the classical dual-pipette assay is replaced by fibronectin-coated polydimethylsiloxane (PDMS) pillars placed perpendicular to the pipette axis and parallel to the imaging plane of an inverted microscope (Fig. 2a and Supplementary Fig. 3a-c). Mature, pre-existing doublets formed over 18 hrs in suspension culture (pre-existing doublets) in ultra-low adhesion dishes at a low cell density, were selected and attached to the side of a pillar tip, and a second doublet was made to adhere to the first doublet in series (Fig. 2a, b and Supplementary Fig. 3d). The cell-cell junction between the two doublets matured over 1 hr; this is an optimal timescale for cells from all groups to form sufficiently strong cell-cell adhesions that cause pillar deflection during cell-cell separation while avoiding cell detachment from the pillar. The two doublets were separated from each other using a pipette by aspirating the free end of the 4-cell complex and manually moving the pipette along the y-axis and away from the pillar. This is performed using a micromanipulator at an average speed of ∼3 µm/sec, while acquiring images at a rate of ∼1 image per second to record the deflection in the pillar (Fig. 2c and Supplementary Fig. 3d). Thereafter, the two cells of the pre-existing doublet still attached to the pillar are similarly separated (Fig. 2d and Supplementary Movie 2). After 1 hr contact and pre-existing (18 hrs) contact SFs were quantified from the respective pillar deflections (Equation-1, see Materials and Methods).

**Figure 2:**
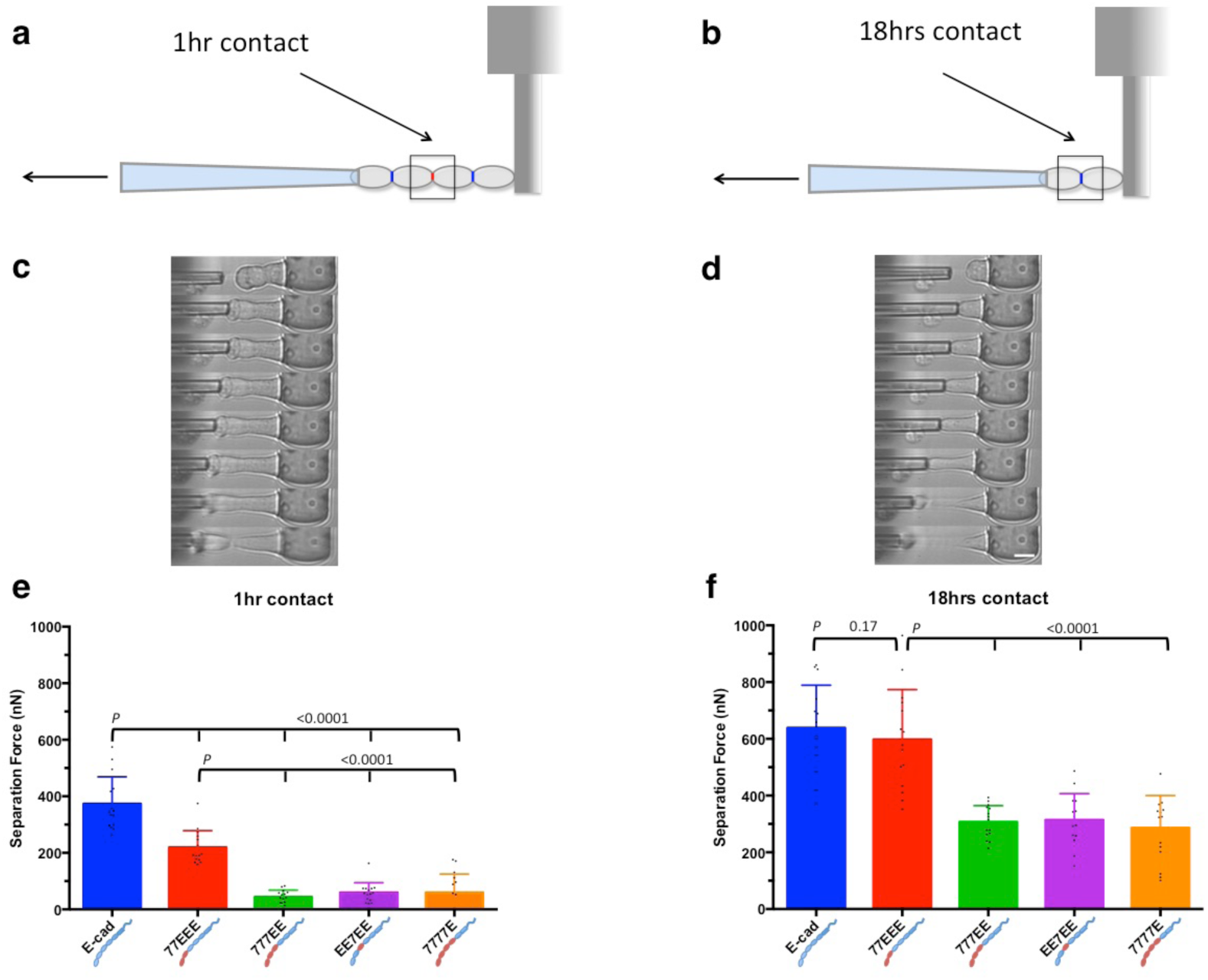
Pillar-pipette assay. **(a)** Cartoon showing two pairs of pre-existing (18 hrs) doublets adhering to each other through a junction established after 1 hr (red, shown by the square box). The doublets are attached to a pillar tip and are pulled by a micropipette to the left. **(b)** Once the two pre-existing doublets are separated from each other, the two cells of the pre-existing doublet with the intact junction (blue, shown by the box) still attached to the pillar are separated. **(c)** Representative kymograph from E-cad group showing two pre-existing doublets adhering to each other through a 1-h contact, and the 4-cell complex attached to a pillar tip. Note the deflection in pillars as the cells are pulled by the pipette. **(d)** Representative kymograph from E-cad group showing a pre-existing doublet with a mature contact and attached to a pillar tip. Scale bar, 15 µm. **(e)** Histogram showing the separation force (SF) for cells expressing wild-type E-cad and chimeric cadherins at 1 hr cell-cell contact (*n* = 15 each). **(f)** Histogram showing the SF for pre-existing (18 hrs) doublets expressing wild-type E-cad and chimeric cadherins (E-cad, *n* = 15; 77EEE, *n* = 15; 777EE, *n* = 14; EE7EE, *n* = 14; 7777E, *n* = 15).

T short times (1hour), the SF for all chimeras missing the EEC3 (777EE, EE7EE, 7777E) was reduced by 8 fold compared to wild-type E-cad after both 1 hr and 18 hrs contact (*P <* 0.0001) (Fig. 2e,f and Supplementary Movie 2). In contrast, 77EEE junctions displayed a 2 fold reduction only. It suggests that EEC3 is essential for stronger SF short-term contacts.

At longer time (18h) the SF strengthened for every chimeric cadherins. 777EE, EE7EE, 7777E only displayed a 2 fold reduction in SF (*P* = 0.17). 77EEE was indistinguishable from wild type E-cad (*P* < 0.0001). It suggested SF matured faster by strand-swapping in EEC1 than in 7EC1, provided that EEC3 is present. To confirm that the differences in SF observed here did not result from differences in cadherin surface levels, we normalized the SFs to cadherin surface levels. Normalized SFs at both time points followed the same trend as the raw SFs (Supplementary Fig. 4a, b), thus data normalization to cadherin surface levels was considered dispensable from now on. In sum, the EEC3 is essential for strong cell-cell SFs, and together with EEC1,2 for fast strengthening.

### EEC1-3 and EEC4 antagonistically modulate single cell cortical tension

In the cell-cell SF assay, we noticed that cells expressing wild-type E-cad and 7777E showed larger deformations during separation as compared with 77EEE, 777EE and EE7EE (Supplementary Movie 2). Since cell cortical tension may affect cell deformation during this assay, and that intercellular adhesion strength depends on cell cortical tension^28,29^, we reasoned that different cadherin EC domains may have distinct effects on cortical tension.

To test this, we used the micropipette aspiration technique (Fig. 3a,b) to characterize the cortical tension of single cells and doublets derived from confluent 2D cultures by trypsin-less dissociation. The cortical tensions of cells expressing chimeric cadherins 77EEE, 777EE and EE7EE were significantly higher than that of cells expressing the wild-type E-cad or the chimeric 7777E (Fig. 3c, *P* < 0.0001). The cortical tension of parental S180 cells (devoid of E-cad expression) was significantly lower than that of E-cad–expressing cells. Finally, the difference in cortical tension between E-cad and 7777E cells was much smaller than between these three groups and other chimera-expressing cells. Strikingly, these results show that changing a single, specific EC domain in an intercellular adhesion protein appears to be sufficient to induce changes in single cell mechanical property.

**Figure 3:**
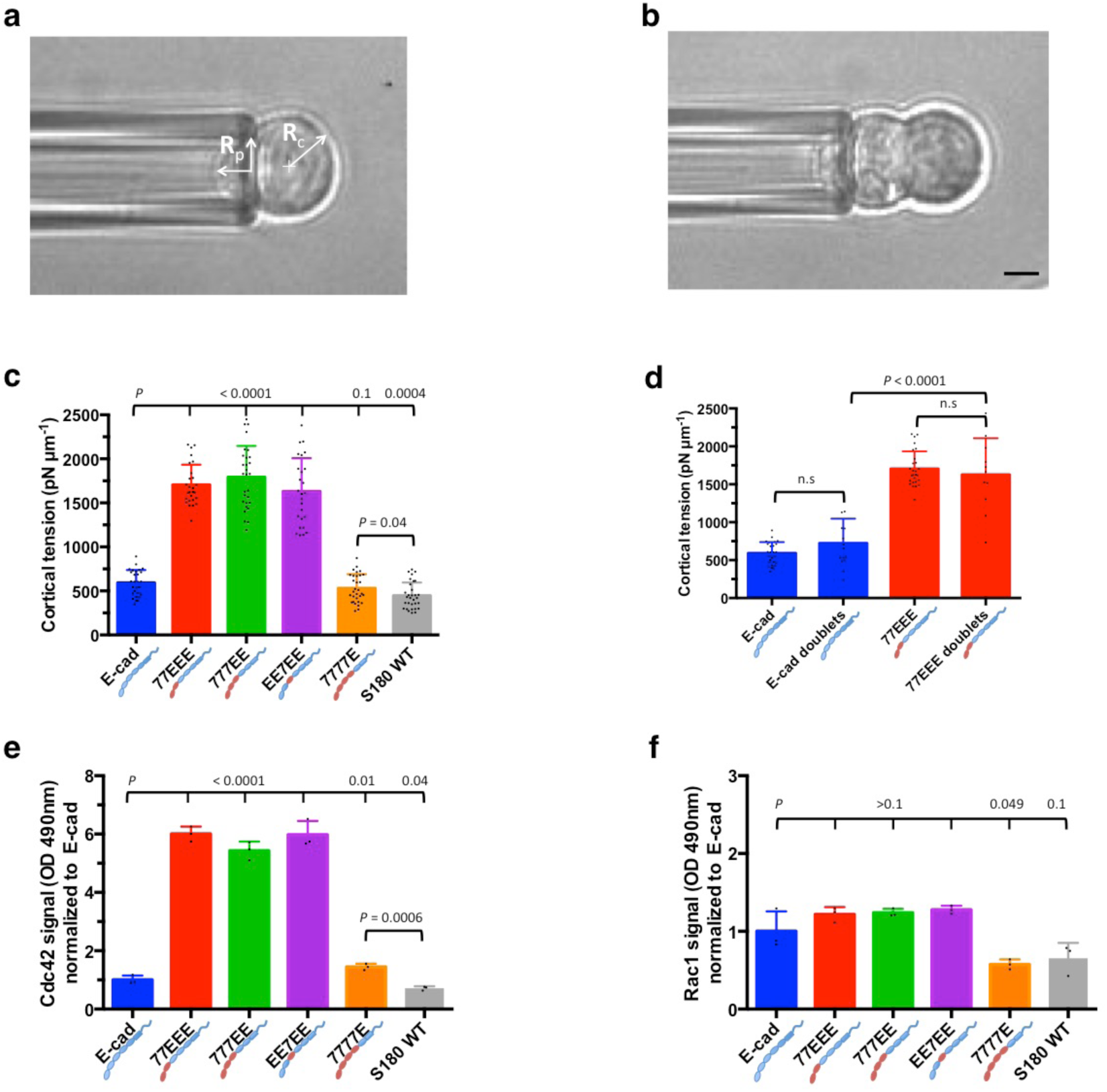
Cell cortical tension and Rho GTPase activity. **(a)** Bright-field image of a single cell held at the pipette by reducing the pressure inside the pipette such that the length of protrusion of the cell inside the pipette is equal to the radius of the pipette R_p_. R_c_ is the radius of the cell. Scale bar, 5 µm. **(b)** Bright-field image of a cell doublet held at the pipette by reducing the pressure inside the pipette such that the length of protrusion of the cell inside the pipette is equal to the radius of the pipette R_p_. Scale bar, 5 µm. **(c)** Histogram of cortical tension values of cells from all groups and the parental S180 wild-type (WT) cells with no endogenous cadherins. Note that cortical tensions of E-cad- and 7777E-expressing cells are significantly higher than that of parental S180 WT cells; cortical tensions of 77EEE-, 777EE- and EE7EE-expressing cells are about three times higher than the parental S180 WT cells. (E-cad, *n* = 28; 77EEE, *n* = 27; 777EE, *n* = 29; EE7EE, *n* = 26; 7777E, *n* = 31; S180 WT, n = 31). **(d)** Histogram comparing cortical tension of single cells and doublets of E-cad- and 77EEE-expressing cells. Following the trend of single cells, the cortical tension of 77EEE-expressing doublets is significantly higher than that of E-cad-expressing doublets. Note that the difference in cortical tension between single cells and doublets is not significant (n.s.) within the same group. However, the spread of values is wider for doublets (E-cad, *n* = 15; 77EEE, *n* = 11). **(e)** Histogram of active Cdc42 signals normalized to E-cad group, showing a 5- to 6-fold higher level of active Cdc42 in 77EEE, 777EE and EE7EE groups. The active Cdc42 in E-cad-expressing cells is significantly higher than that of the parental S180 cells but significantly lower than that of the 7777E-expressing group (*n* = 3). **(f)** Histogram showing active Rac1 levels normalized to the E-cad group. Note that the active Rac1 levels in the 7777E group were marginally lower than those in the E-cad group. The levels in 77EEE, 777EE and EE7EE groups were higher than that in the E-cad group but not statistically significant (*n* = 3).

To assess whether cortical tension also depends on intercellular adhesion, we then measured differences in cortical tension in doublets of E-cad- or 77EEE-expressing cells. We observed no significant difference between cortical tension of single cells and doublets of the same group (*P =* 0.07 and *P =* 0.5 respectively), and the difference in cortical tension between E-cad and 77EEE doublets was significant (Fig. 3d, *P* < 0.0001). Therefore, cortical tension depends on the EC regions of expressed cadherins but not on intercellular adhesion.

### Cadherin EC region modulates GTPase signalling to alter cortical tension

To investigate how cadherin EC region may affect cell cortical tension, we examined the activity of Cdc42 and Rac1 using a G-LISA assay on cell lysates. Indeed, distinct Rac1 activity—both cell-autonomous and cadherin *trans*-bond-dependent—occurs in response to the expression of different cadherin proteins in the same cellular background^35^. When compared with E-cad–expressing cells, we found a substantial increases in the active Cdc42 levels in cells expressing 77EEE (*P* < 0.0001; Fig. 3e), 777EE (*P* < 0.0001), EE7EE (*P* < 0.0001). Cells expressing 7777E also exhibited an increase (*P* = 0.01), albeit much smaller. Active Cdc42 levels were also low in the parental S180 cells (*P* < 0.04). Active Rac1 levels of all the cell lines, however, were not significantly different from that of the E-cad expressing cells (*P* ≥ 0.1), with the exception of 7777E-expressing cells, which was lower (*P* = 0.049) (Fig. 3f). These results show that Cdc42 activity levels correlate with cell cortical tension, and therefore suggest that the EEC and 7EC regions may differentially regulate cortical tension through Cdc42.

### Cell-cell separation forces are governed by dissipative processes

Within a previously established theoretical framework^8^, it is possible to assess the contribution of static intercellular adhesion energy to cell-cell SF. Indeed, the static intercellular adhesion energy W is defined as W = γ(1-cosθ), where γ is the cortical tension and cosθ the compaction parameter (Fig. 4a). In turn, the static SF, SF_static_ = πRW, where R is the cell radius, in the limit of low compaction. Finally, the contact radius r is determined by r = Rsinθ. Thus, we set out to measure cell-cell compaction and the contact radius.

**Figure 4:**
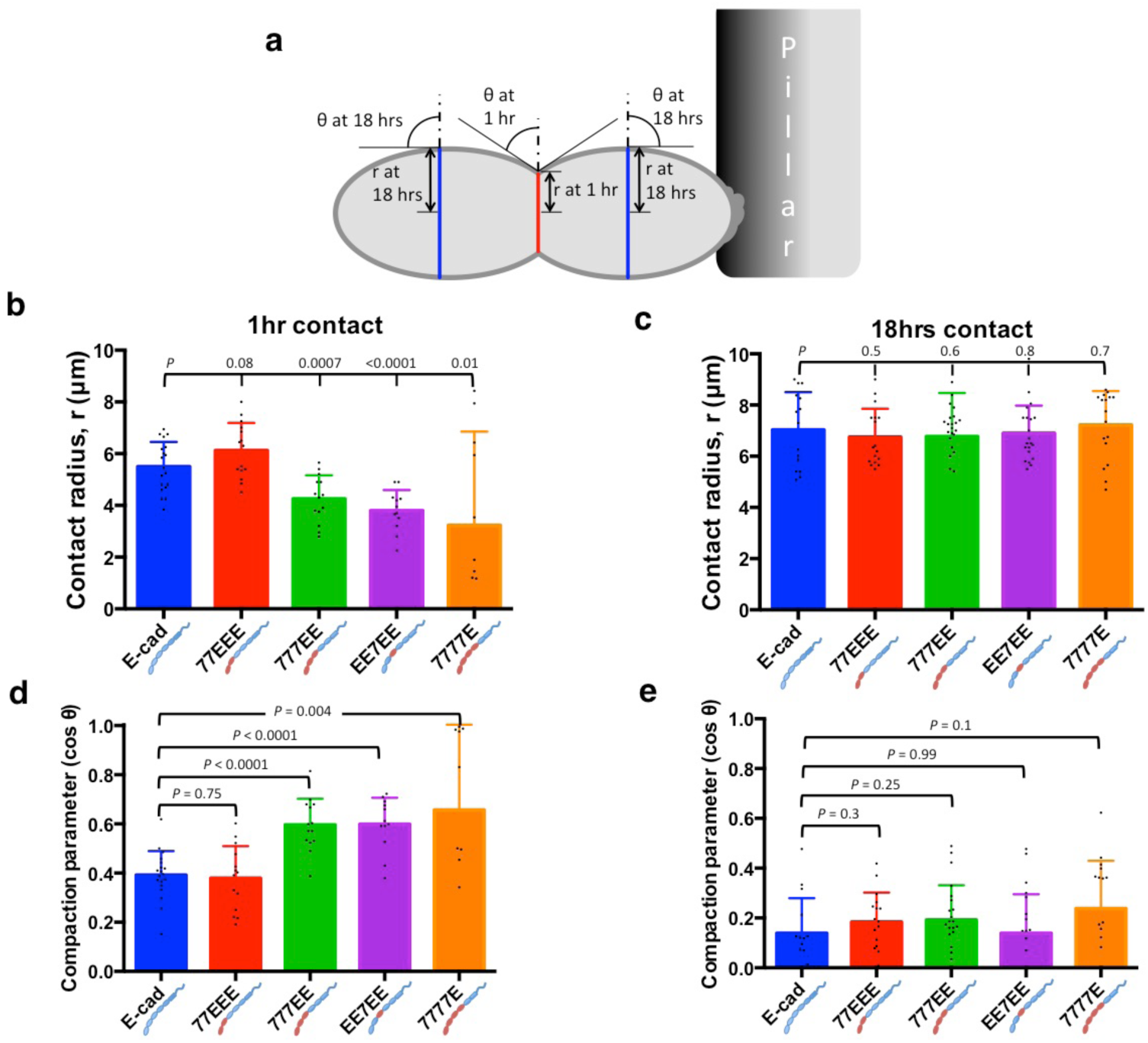
Contact radius and compaction parameter. **(a)** Drawing depicting two pre-existing (18 hrs) doublets adhering to each other through a 1-h contact. The 1-h cell-cell junction is shown in red and the pre-existing (18-h) junctions in blue. Measurements of the cell-cell contact angle “θ” and the contact radius “r” in 1 hr and pre-existing (18 hrs) doublets are illustrated. **(b)** Histogram of contact radius at 1 hr contact (E-cad, *n* = 19; 77EEE, *n* = 15; 777EE, *n* = 14; EE7EE, *n* = 12; 7777E, *n* = 10). **(c)** Histogram of contact radius in pre-existing (18 hrs) doublets (E-cad, *n* = 15; 77EEE, *n* = 17; 777EE, *n* = 23; EE7EE, *n* = 21; 7777E, *n* = 17). Note that the mean contact radii of all groups are similar, although their separation forces (SFs) are significantly different (Fig. 2f, h). **(d)** Histogram of dimensionless compaction parameter (cos θ) at 1 hr of contact (E-cad, *n* = 19; 77EEE, *n* = 14; 777EE, *n* = 14; EE7EE, *n* = 12; 7777E, *n* = 10). Note that, although the compaction parameter of E-cad and 77EEE groups are similar, their SFs are significantly different, as seen in Fig. 2e, g. **(e)** Histogram of compaction parameters of pre-existing (18 hrs) doublets (E-cad, *n* = 15; 77EEE, *n* = 17; 777EE, *n* = 23; EE7EE, *n* = 21; 7777E, *n* = 17). Note that the compaction parameters for E-cad and other groups are similar, even though they show significantly different SFs (Fig. 2f, h).

At 1 hr of contact, the trends of the mean contact radius and compaction parameter were seemingly indicative of the measured SF across different groups (Fig. 4b,d). Indeed, E-cad- and 77EEE-expressing cells showed insignificant differences in contact radii and compaction parameters that were higher and lower, respectively, than that of the other chimeric EC cells. The SF, r and cosθ metrics appeared to behave consistently with higher intercellular adhesion energy in E-cad- and 77EEE-cells compared with other chimeric EC cells. After 18 hrs, however, both the contact radius and compaction parameter narrowed to around 7 µm and 0.18, respectively, for all groups, without significant differences between groups (Fig. 4 c, e).

We next computed the SF_static_ from cortical tension and compaction parameters at 1 hr (where low compaction is a reasonable assumption) and 18 hrs. We found that SF_static_ scaled well below the measured SFs in all conditions and, notably, at least an order of magnitude below that measured in cells expressing E-cad or 77EEE (Table 1). We conclude that static adhesion energy only marginally contributes to cell-cell SFs and that EEC1-3 contributions to cell-cell SFs essentially arise from dissipative processes.

**Table 1:** Comparison between measured separation force and computed static separation force (SF vs SF_Static_)

|  | Measured |  |  |  |  |  |  |  | Calculated |  |  |  |  |  |
| --- | --- | --- | --- | --- | --- | --- | --- | --- | --- | --- | --- | --- | --- | --- |
| | SF 1h, (nN) | SF 18 h (nN) | R 1h, ($\mu\text{m}$) | R 18h, ($\mu\text{m}$) | $\cos\theta$ 1h | $\cos\theta$ 18h | $\gamma$ 1h, (pN/$\mu\text{m}$) | $\gamma$ 18h, (pN/$\mu\text{m}$) | $R=r/\sqrt{1-\cos^2\theta}$ 1h | $R=r/\sqrt{1-\cos^2\theta}$ 18h | $W=\gamma(1-\cos\theta)$ 1h | $W=\gamma(1-\cos\theta)$ 18h | $SF_s = \pi RW$ 1h (nN) | $SF_s = \pi RW$ 18h (nN) |
| E_cad | 373 | 639 | 5.5 | 7 | 0.4 | 0.15 | 590 | 722 | 6.0 | 7.080 | 354 | 613 | 6.7 | 13.7 |
| 77EEE | 218 | 598 | 6 | 7 | 0.4 | 0.18 | 2 | 2 | 6.55 | 7.12 | 1021 | 1334 | 21.0 | 29.8 |
| 777EE | 44 | 307 | 4.5 | 7 | 0.6 | 0.18 | 2 |  | 5.62 | 7.12 | 716 |  | 12.7 |  |
| EE7EE | 59 | 314 | 4 | 7 | 0.6 | 0.13 | 2 |  | 5.0 | 7.06 | 651 |  | 10.2 |  |
| 7777E | 59 | 286 | 3 | 7 | 0.6 | 0.21 | 529 |  | 3.82 | 7.16 | 201 |  | 2.4 |  |
| S180 |  |  |  |  |  |  | 446 |  |  |  |  |  |  |  |

### EEC3 and EEC1-3 provide tensile strength and ductility, respectively, to epithelial cell sheets

Given the uncorrelated effects of the EC region on cell-cell SFs and cortical tension, we sought to assess how this combination would affect the mechanical properties of epithelial tissues. To avoid the confounding effects of cell-substrate adhesion, we developed a dispase-based epithelial tissue uniaxial stretcher that locally cleaves cell-substrate bonds, causing a local stretch on the adherent cell sheets in culture, and thereby shifting cell-generated tension entirely to cell-cell junctions (Fig. 5a-d).

**Figure 5:**
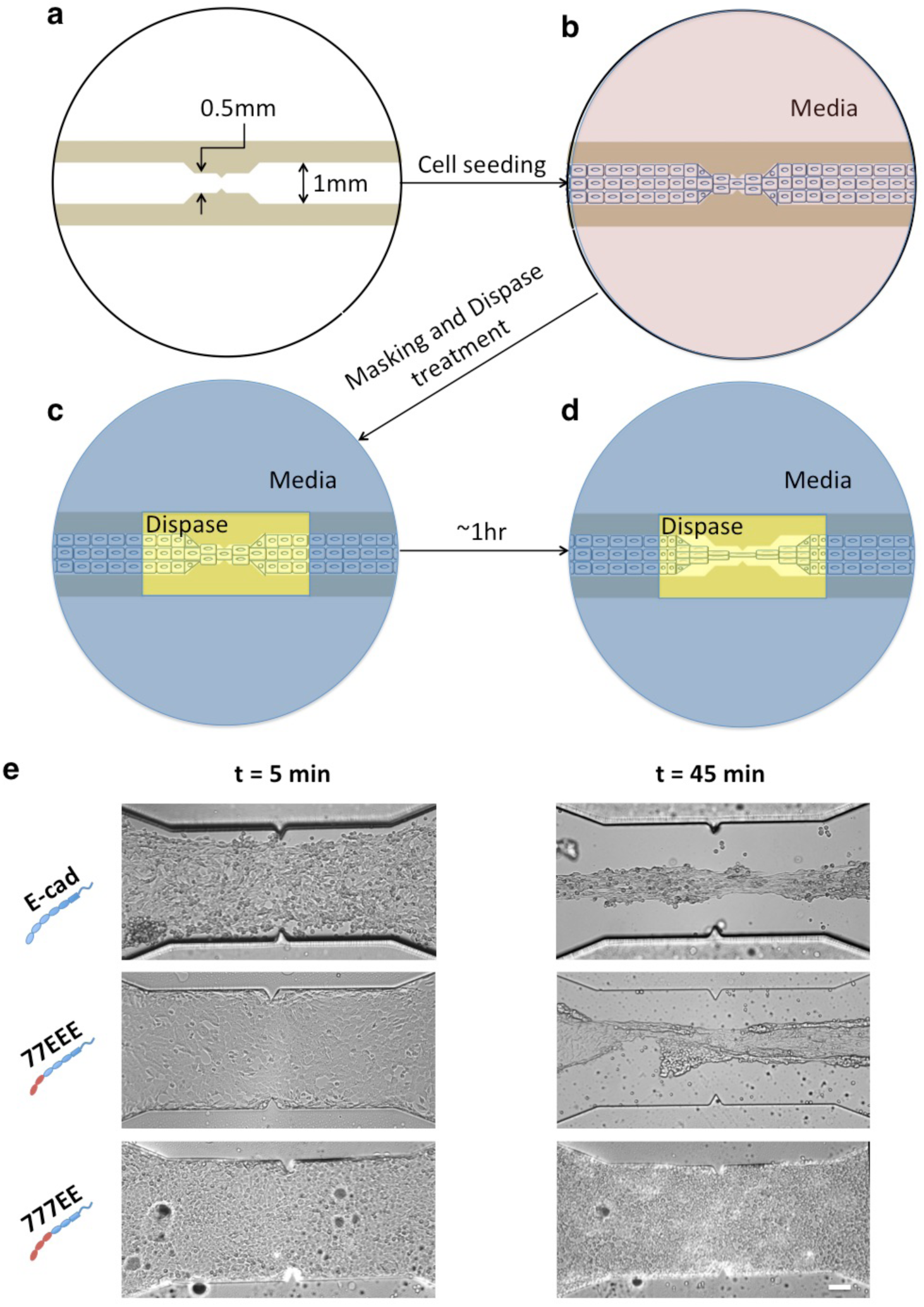
Dispase-based junctional protein tensor assay. **(a)** Schematic of a 1-mm wide channel with a central constriction of 0.5 mm made of UV curable polymer on a 27-mm glass-bottomed dish. **(b)** Channel seeded with cells forming cell-cell junctions. Fewer cells are shown here for ease of representation. **(c)** PDMS mask (blue) with a 5 mm × 2.5 mm opening in the middle that snuggly fits inside the dish. The media is removed from the opening and dispase (yellow) is added. The media remains in the area covered by the PDMS mask. **(d)** After ∼1 hr of dispase treatment, cell-substrate bonds within the area are cleaved while the cells under the PDMS, where the culture medium is present, are still attached to the substrate. In the absence of cell-substrate adhesion, the cells that become lifted in the constricted dispase-treated region experience tension and stretch in the direction along the channel at the expense of cells contracting at either side of the constriction due to the intrinsic cortical tension of the cells. Figure not to scale. **(e)** E-cad-, 77EEE-, and 777EE-expressing S180 cells subjected to dispase treatment. Imaging was commenced 5 min after the addition of dispase for 1 hr. Note that, in the t = 45 min image, the E-cad-expressing cells have undergone significant strain yet the cell-cell contacts are still maintained. In the 77EEE-expressing cells, the cell sheet snaps when the sheet is lifted at lower strain levels as compared to E-cad-expressing cells. Although cell-cell junctions are formed similarly in 777EE- and 77EEE-expressing cells, 777EE-expressing cells are not able to maintain cell-cell contact in the absence of cell-substrate adhesion; they became single cells and cell clumps that are unable to withstand the tension as a cell sheet. Scale bar, 100 µm.

We found that, following dispase treatment, the E-cad cell sheet could withstand cell-generated tension while thinning as a ductile material (Fig. 5e and Supplementary Movie 3). This was accompanied by cell reorientation along the direction of tension. Comparatively, the initially cohesive 77EEE cell sheet soon snapped under dispase treatment—even though the two parts of the sheet each remained cohesive upon tension relief (Fig. 5e and Supplementary Movie 3)—whereas the 777EE and EE7EE cell sheets disintegrated to single cells and cell clumps with tethers in between, respectively (Fig. 5e and Supplementary Movies 3&4). Finally, cells expressing 7777E failed to form cohesive cell sheets (Supplementary Movie 5), consistently with the inability of this chimera to support epithelial integrity (see above). Considering ductility as the inverse of the sheet cross-section area at rupture and tensile strength the pulling force at rupture, these results suggest that EEC3 alone provides tensile strength to the cohesive cell sheet, while together with EEC1,2 it provides ductility, so that the sheet can withstand larger strains before rupture

Since the main phenotypic differences between wild-type and 77EEE cells appeared at the tissue level, in terms of ductility, we sought to determine the relevance of EC1,2-mediated dimerization in this process. As we had previously shown that epithelial integrity was impossible without strand-swapping (Supplementary movie 1, left panel), we stably expressed X-dimer incompetent (K14E mutation) full-length E-cad mutants tagged with eGFP in S180 cells by lentivirus-based transduction. In contrast with cells expressing strand-swap mutants, which bear no tensile-strength, cells expressing X-dimer-incompetent E-cad formed stable junctions comparable to those formed by cells expressing wild-type E-cad, and showed a ductile behavior similar to that of wild-type E-cad expressing cells (Supplementary movie 6). Thus, these results show that X-dimerization is neither essential nor accountable for differences in tissue ductility, pointing to a cell-autonomous mechanism, as for cell cortical tension.

### EEC1-3 are essential for collective cell migration speed

A study proposed how cell-cell adhesion and cell cortical tension governed the dynamics of collective behaviours in adherent cell assemblies^36^. Thus, we next sought to assess how altering the material properties of cell sheets with different ECs would impact dynamic multicellular processes using wound-free gap closure assays.

Cells expressing 7777E showed the fastest gap closure compared with the other EC chimera-expressing cells (Fig. 6a-c and Supplementary Movie 7) but slower than E-cad expressing cells (*P* = 0.008), and this occurred as a non-cohesive cell assembly. This is consistent with epithelial integrity impeding cell migration speed. Nevertheless, the rate of gap closure for E-cad-expressing cells was twice as fast as that of 77EEE-expressing cells (*P* < 0.0001); while the rate of 777EE- and EE7EE-expressing cells was significantly lower than that of E-cad (*P* < 0.0001). (Fig. 6c). As the EEC3 seems to aid faster migration and the lack of EEC4 abolishes cell cohesivity during migration, we wanted to check if just EEC3 and EEC4 are sufficient for fast and collective cell migration. We designed a 77EE7 chimera with the transmembrane and cytoplasmic domains from cad-7 and expressed in S180 cells. Owing to a low expression level, (Supplementary fig. 5a), the mean SF of 77EE7 doublets at 18 hrs of contact was also low (147 ± 58 nN) and comparable to the mean SF of 7777E expressing cells (*P* > 0.2) (Supplementary fig. 5b). However, despite low SF, the 77EE7 expressing cells migrated collectively and at a higher speed than the 777EE cells (Supplementary Fig 5c). Together, these results are consistent with a positive effect of EEC3 on collective cell migration, which provides higher tensile strength, and of EEC1-3, which provides higher ductility, in a manner that compensates epithelial integrity effects caused by EEC4. Remarkably, this verifies previous predictions^36^.

**Figure 6:**
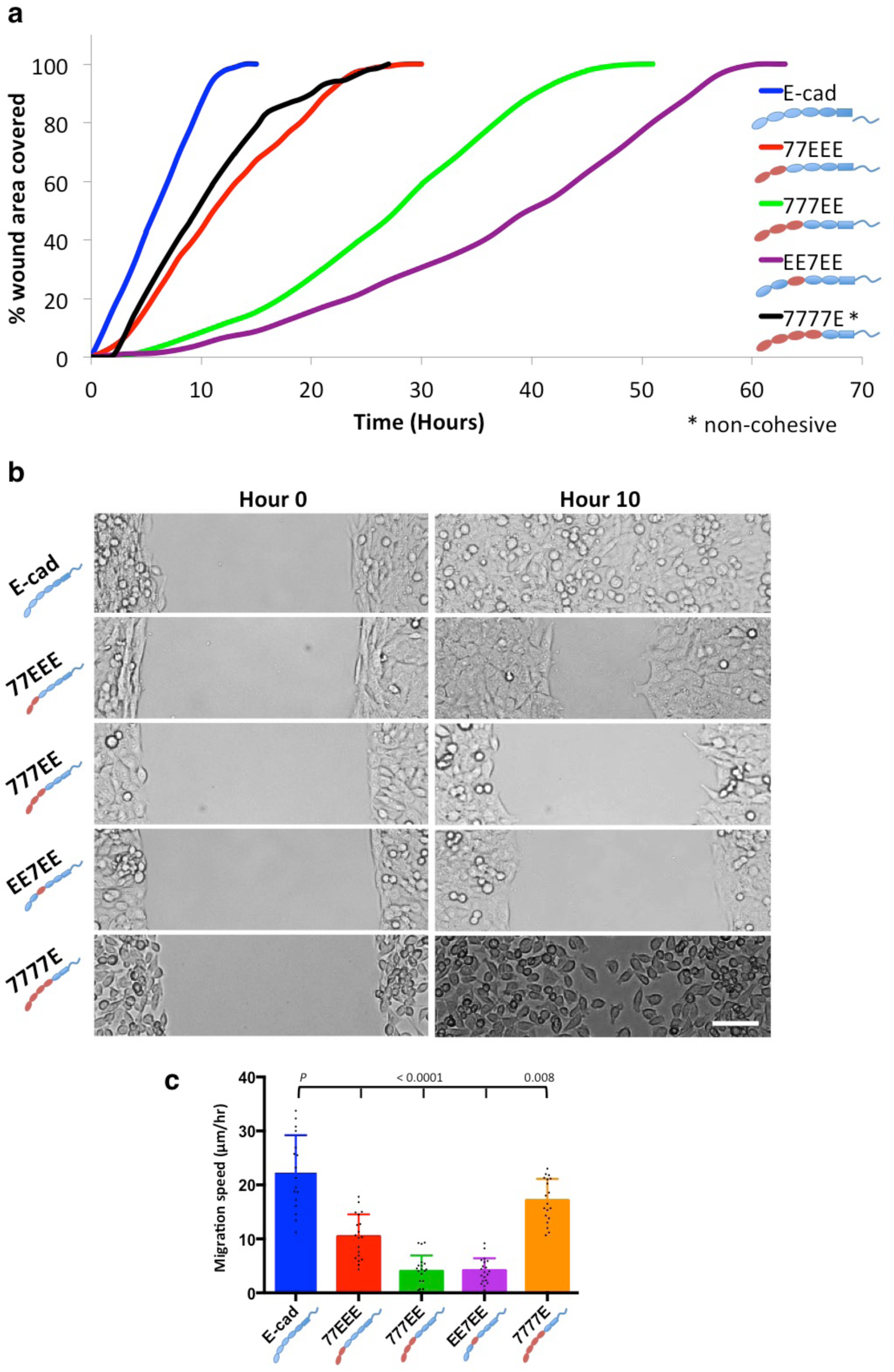
Wound healing assay. **(a)** Representative curves showing the time taken to cover a 500-µm unwounded gap. All groups migrated collectively, except 7777E-expressing cells. E-cad expressing cells migrated fastest, followed by 77EEE-expressing cells. 777EE- and EE7EE-expressing cells took approximately 4- to 5-times longer than the time taken by E-cad-expressing cells. The migration speed of 7777E-expressing cells is comparable to that of E-cad-expressing cells; however, these cells migrated as a non-cohesive cell assembly, as the cell-cell adhesion in these cells is transient. **(b)** Representative images from each group showing the position of cells at the beginning and end of 10 h of migration. Scale bar, 50 µm. **(c)** Histogram showing the cell migration speed of all the groups. (E-cad, *n* = 17; 77EEE, *n* = 19; 777EE, *n* = 20; EE7EE, *n* = 20; 7777E, *n* = 19).

A summary of the results is presented in Table 2.

**Table 2:** Summary of results. We classified the influence of the different chimeric Cadherins on the selected phenotypes as *** High, ** Moderate and * Low.

| Phenotypes \ Cadherins | E-cad | 77EEE | 777EE | EE7EE | 7777E |
| --- | --- | --- | --- | --- | --- |
| Cell-cell adhesion strength | *** | ** | * | * | * |
| Cortical tension | * | *** | *** | *** | * |
| Collectivity / colony formation | *** | *** | *** | *** | * |
| Speed of migration/motility | *** | ** | * | * | *** |

## Discussion

Cadherin expression is an important determinant of intercellular adhesive properties^6^. Here we sought to characterize how Sthe EC domains of type-I and type-II cadherins affect cell-cell adhesion and tissue phenotypes. We expressed chimeras of E-cad and cad-7—preserving the calcium binding, transmembrane, and cytoplasmic regions of E-cad but swapping the various ectodomains of the two proteins—in cadherin-null cells, and sorted for medium overall expression based on eGFP. Although cadherin surface levels were not identical among the chimera-expressing cell groups (possibly due to small differences in expression levels), junctional cadherin levels could be modulated by the cadherin-type of EC domains. Indeed, EEC1-3 but not EEC1-4 were dispensable to accumulate cadherins at cell-cell contacts and recruit catenins and vinculin, thereby promoting cytoskeleton anchoring and adhesion strengthening^37^,^38^. This shows that EC domains can regulate the recruitment of cadherin cytoplasmic partners through the control of cadherin incorporation at cell-cell contacts, in a cadherin-type specific manner. However, this cadherin-type specific recruitment of cytoplasmic partners does not rely on the strand-swap mechanism alone, although strand-swapping is required (as cells expressing strand-swap incompetent E-cad mutants didn’t form stable junctions). Instead, our results are consistent with a role for EEC4 in this process, since EEC5 ectodomain is insufficient for epithelial integrity. Interestingly, hypoglycosylation of EEC4, which occurs upon densification of cultures of epithelial cells, impacts the composition and stability of the cadherin/catenin complex, and impedes collective cell migration^39,40^. Moreover, normal glycosylation of EC4 of type-II VE-cadherin was previously found to prevent cadherin cis-clustering^41^. Future studies may investigate whether glycosylation-dependent cis-clustering and/or catenin recruitment mechanisms explains the difference between EC4s of type-I and -II cadherins in our experiments.

We used a pillar-pipette assay to quantify intercellular SFs, as this alleviates some of the potential limitations of the classical dual-pipette SF assay. In the dual-pipette assay, the cortical cytoskeleton is remodeled due to aspiration by the pipettes. Moreover, the dual-pipette assay requires discrete increments in pipette aspiration, resulting in stress-relaxation cycles, which may reinforce the junction^42^, or change the cortical tension, and thus affect the measurements. Previous protocols for preparing pre-existing doublets from cell suspensions for SF assays were based the mechanical dissociation of confluent, 18 hrs cultures by pipetting^26^, but it is very rare to get pre-existing doublets in weakly adhering cell types as the cell-cell junctions often rupture during such mechanical dissociation process. Here, we used suspension cultures seeded at low density on ultra-low adhesion dishes to let cells form long-term doublets in the absence of cell-substrate adhesion by disrupting cell motility on substrates and thus minimizing the chances of doublets getting separated from each other after initial contact. Moreover, we previously observed that stimulating a cell of the doublet with a fibronectin coated bead for one hour promotes an increase in cell-cell SF43. The attachment of cell to the fibronectin coated pillar might also trigger such an increase in cell-cell separation force. These could be the reasons why we get higher SF values for pre-existing doublets as compared with results in previous studies (SF of E-cad: 638 ± 150 nN vs 350 nN).

Intercellular SFs between E-cad-expressing cells are much higher than that between cad-7 expressing cells, and this is mostly due to differences in extracellular domains^6^. Here, our results show that SFs are high only in cells expressing cadherins bearing the EEC3. Previous studies on type-I N-cadherin support that EC3 glycosylation impedes cis-dimerization and thereby affects cadherin *trans*-bond kinetics^44,45^. As for EC4, a differential regulation of EC3 glycosylation-dependent clustering between E-cad and cad-7 may underlie the considerable differences in SFs between these cadherin types.

Previous structural and single-molecule studies have shown that both type-I and type-II cadherins are competent for both EC1,2-dependent strand-swap and X dimers, and proposed that the differences in the EC1 domain between the two types was sufficient to confer the type-specific activities of cadherins in terms of *in vitro* and *in vivo* cell sorting^9,10^. Here, strand-swap or X dimerization specifics of type-I and -II cadherins appear to only transiently affect SFs: SF strengthening being faster between EEC1,2 than between 7EC1-2, perhaps because strand-swapping kinetics differ. Indeed, this difference only arises when EEC3 is present, as if strand-swapping differences were EC3-dependent, reminiscently of EC3 glycosylation effects on *trans*-bond kinetics^45^. In contrast, we show that mature SFs reach similar levels regardless of the type of EC1,2. Thus, differences in mature SFs are unlikely governed by differences in strand-swap or X-dimer properties between cadherin types, and SFs may not predict cell sorting or vice versa. In fact, our quantitative analysis of cell cortical tension and compaction revealed that the static intercellular adhesion energy—the quantity that does govern cell sorting^28,29^—only marginally contributes to cell-cell SFs. Therefore, we speculate that EEC3, possibly through glycosylation-dependent cis-clustering, contributes to dissipative mechanisms that allow cells to mechanically discriminate cell-cell separation from cell sorting. Consistently, we show that EEC3 provides tensile strength to epithelial sheets, which essentially relies on the ability of cells to bear stronger tensile forces before cell-cell contact rupture. Nevertheless, up to half the strength of mature separation forces remains EC type-independent. Thus, the cells appear to achieve with time a cadherin organization within the contact that is sufficient for a substantial contribution of dissipation upon cell-cell separation, regardless of the EC region specifics. This contribution may merely result from the cadherin tail/cytoskeleton driven accumulation of cadherins at the contact, since cadherin tail and cytoskeleton anchoring remains essential for SF strengthening, regardless of cadherin type^6,26^.

While EC-mediated cis-interactions and, at least partly independently, the cytoskeleton may synergize to accumulate stable cadherin complexes at cell-cell contacts, cell-cell separation ultimately involves the rupture of these cadherin-mediated *trans*-bonds. Thus, upon cell-cell separation, a likely contribution to dissipation may merely be cadherin *trans*-bond rupture. Indeed, the energy per unit area G associated with the rupture of flexible bonds between two surfaces is reflected by (n_0_/2k)(k_B_T/Δ) ^2^ln^2^ (v/v_0_), where n_0_ is the bond surface density, k the molecular stiffness of the bond, k_B_ the Boltzmann constant, T the temperature, Δ the interaction spatial range of the bond, v the pulling velocity, and v_0_ the thermal velocity of bond rupture^46^. Using typical values^47^ of n_0_∼10^16^ m^-2^, k∼10^−3^ N/m, Δ∼0.1 nm, v∼3 µm/s (as in our conditions) and v_0_∼0.01 µm/s, G is about 60 mN/m. Scaled to the size R∼5 µm of the cell, this gives a contribution to the SF easily reaching several hundred nN, which is in remarkable accordance with our measured values. This does not exclude that additional bond ruptures between cadherin complexes and the cytoskeleton, or within the cortex may contribute too.

We have previously shown that Rho-like small GTPases, Cdc42 and Rac, are activated in cell aggregates expressing E-cad and play a critical role in the maturation of strong SFs^26^. The activities of Rho GTPases are spatially regulated upon cell-cell contact formation^50^. Pathways involved may implicate aPKC, Par3-Par6, or Elmo2 and Dock1, and result in a local dissolution of the actomyosin cortex^51,52^, which will in turn contribute to the intercellular adhesion energy and promote compaction^28,29^. Interestingly, cadherin-*trans*-bonding, in a cadherin sub-type-dependent manner, regulates Rho GTPase activation and alter cell stiffness as measured by cell twisting cytometry^35^. This supports that cadherin regulation of Rho GTPases affects the actomyosin cortex at a distance from cell-cell contacts in a type-dependent manner. Here, we show in addition that cells exhibit correlated Cdc42 activities and cortical tensions dependent on the cadherin EC domains but independent of intercellular adhesion. Therefore, cadherin EC domains, specifically EEC1-3 and EEC4, are involved in a cell-autonomous outside-in signaling targeting Rho GTPases and cortical tension. Despite the common involvement of EEC4 in cortical tension and epithelial integrity, the lack of correlation between these two properties as a function of EEC1-3 precludes that epithelial integrity emerges from high cell cortical tension. In contrast, the tissue stretching assay reveals a correlation between low cortical tension and tissue ductility, which supports a model by which tissue ductility arises from cell-autonomous low cortical tension. Along the same line, cells expressing E-cads competent or incompetent for X-dimerization show the same ductile behavior as a tissue, further supporting that cortical tension and ductility—provided that stable cell-cell junctions can occur—are regulated independently of cadherin *trans*-bond regulation, and are therefore cell-autonomous.

A previous theoretical study had showed how cell-autonomous mechanical properties could determine collective behavior of epithelia, specifically implicating cell cortical tension, increase in which impedes epithelial remodeling, akin to a jamming transition^36^. Here, wound-healing assays reveal an apparent complex dependence on cadherin EC domains. Nevertheless, the effects of EC domains on intercellular contact formation, SFs, cortical tension, tissue tensile strength, and tissue ductility allow us to propose the following model. First, the formation of the cell layer devoid of intercellular gaps, supported by EEC4, dampens single-cell migration speed, and prevents the cohesive sheet from migrating as fast as a collection of individual cells, reminiscently of a density-driven jamming^36^. Second, the EEC3 provides tissue tensile strength, which allows cells to pull harder on each other while migrating, thereby promoting collective migration speed. Note that we had previously demonstrated that cadherin-mediated adhesion stimulates cell spreading and that stronger cell-cell adhesion results in stronger traction forces at focal adhesions through the Src and P13K signaling pathways^53^. This mechanism could also contribute to faster collective cell migration speeds in cells with stronger cell-cell adhesion. Finally, EEC1,2 provide tissue ductility, which allows cells to undergo higher strains during collective migration, as if cells unjammed from a cortical-tensioned state^36^, thereby allowing even greater collective migration speed, (Fig. 7). Thus, fast collective migration emerges from a fine balance of tissue tensile strength and ductility.

**Figure 7:**
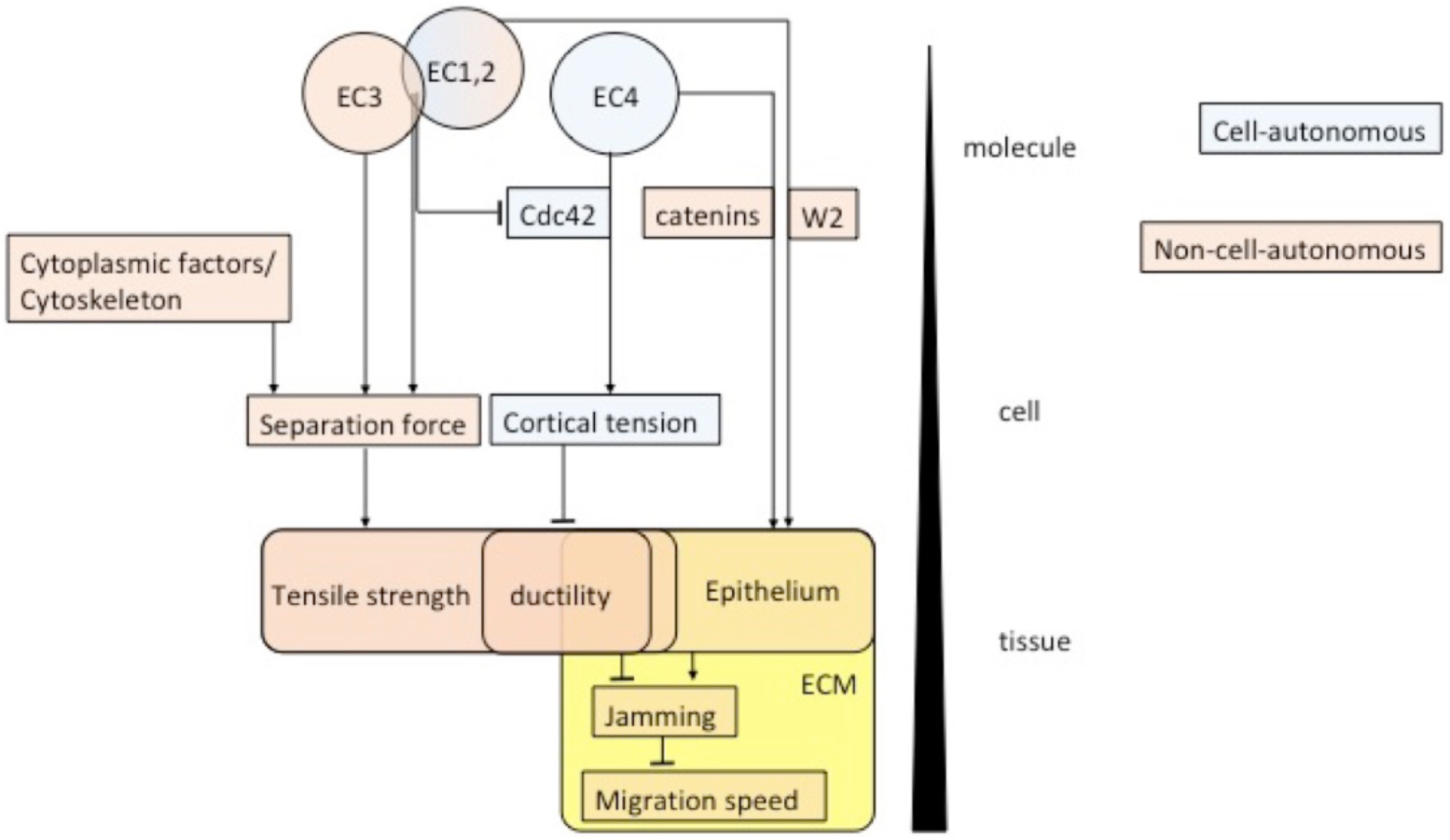
Proposed model depicting the influence of E-cadherin EC1-4 domains on cell-autonomous and non-cell-autonomous cell and tissue properties. EEC4 regulates catenin recruitment and formation of epithelia on the extracellular matrix, and, antagonistically with EEC1,2 but in a Cdc42-dependent manner, cell– autonomous cortical tension. EEC1,2 also with EEC3 regulate intercellular separation forces together with the cytoskeleton and other cytoplasmic factors. High intercellular separation forces support tissue tensile strength while low cortical tension promotes tissue ductility. Tissue ductility in turn promotes intercellular unjamming and thereby antagonizes the dampening effect of EEC4-dependent epithelial phenotype on collective cell migration speed.

In summary, using a range of microscopy, cell and tissue manipulation assays, we determined that the EEC4 region promotes the recruitment of catenins and vinculin to cell-cell contacts and epithelium-like cohesive cell sheet formation on an extracellular matrix. EEC4 also contributes to the increase in cell cortical tension in an intercellular adhesion-independent manner, likely through a Cdc42 pathway. In turn, the EEC3 is required to provide strong cell-cell SF and, subsequently, tensile strength of the cell sheet, mostly through an increase in dissipation during cell-cell separation rather than an increase in static intercellular adhesion energy. Moreover, EC1,2 together with EEC3 provide an additional increase in cell-cell SF and tensile strength but also antagonize EC4 activity on cell cortical tension for increased ductility to the cell sheet. Finally, we propose that increased ductility compensates the dampening effect of epithelium formation on cell migration speed. Cortical cytoskeleton regulation by the cadherin cytoplasmic region, rather than cadherin *trans*-interactions, has increasingly appeared as a key determinant of intercellular adhesion energy and cell sorting. Our findings nevertheless remind us of the importance of cadherin extracellular domains in major tissue mechanical properties, and reveal their respective effects and underlying cellular mechanisms. Whether the expression of full-length type-I and type-II cadherins in varying proportions in the same cells throughout development determines the major morphogenetic events through the mechanisms unveiled here is a promising research avenue for future studies.

## Compteing interests

The authors declare no competing interests for the work described in this manuscript

## Methods

### Cell lines, constructs, lentivirus-based transductions, and immunostaining

Protein sequences and domain organizations of mouse E-cad and cadherin-7 were used to construct chimeric cadherins, including 77EEE, 777EE, 7777E and EE7EE, as shown in Fig. 1a, b. In these chimeras, we preserved the calcium binding sites. Sequences were cloned into PLVX-puro vector with eGFP at the C-terminus and proper insertion confirmed by sequencing. HEK293T cells were co-transfected (Effectene transfection reagent) with the plasmid and optimized lentiviral packaging mix (ViraPower™). The viral supernatant was harvested, filtered, and added to S180 cells plated at 80% confluence. Stably transduced cells were selected based on puromycin resistance, FACS sorted for a medium level of expression, and checked for mycoplasma contamination (MycoAlert™, Lonza). Rabbit anti-α-catenin (Sigma-Aldrich), mouse monoclonal anti-β-catenin, mouse anti-p120-catenin (Invitrogen) and mouse monoclonal anti-vinculin (Sigma-Aldrich) were used for immunostaining. Strand-swap incompetent E-cad (W2A mutant) and X-dimer incompetent E-cad (K14E) were also stably expressed separately in S180 cells following the above lentivirus-based transduction protocol and FACS sorted for medium expression.

### Computer model for validation of preserved domain interactions and calcium binding sites in chimeric cadherin ECs

Models of juxtaposed domains at the hybrid interfaces were built using the web-based interface of SWISS-MODEL^34^ and existing protein structures as templates: 2A62, 3Q2V, 3Q2W, 1L3W and 5VEB^5,9,47,54^. Calcium ions were placed by hand and sidechains angles were orientated to coordinate the ions in Coot^55^.

### Confocal microscopy, image processing and analysis

Confocal images on fixed samples were acquired using a Nikon A1plus microscope. Z-stacks of cells were acquired from 1 µm below the basal cell surface to 1 µm above the apical cell surface, with a step size of 0.5 µm at 60x magnification. The laser power, pinhole size and other standard parameters were kept identical for all samples. ImageJ software was used for quantification of signal intensities. For quantifying the cadherin cell surface intensity, the image slice containing the planar cell surface in contact with the coverslip was detected and one slice above and below the identified image were taken. A maximum intensity z projection was performed on the three images. Using the freehand line tool, with a line width of 3, a line of approximately 20 – 30 µm long was drawn in the region between the cell edge and the nucleus. Using the plot profile function, the intensity plot was obtained and the intensity values were listed and saved in Excel sheets for data processing. In a similar manner, data was collected from the cell free area of the coverslip and subtracted to eliminate the background signal. The cell surface cadherin signal intensity per µm length was calculated and normalized to E-cad per µm intensity. In a similar manner, junctional cadherin intensity was quantified by selecting all the slices in the stack for maximum intensity z projection and tracing the cell-cell junctions with the freehand line tool.

### Tissue culture, cell dissociation

Cells were maintained in high glucose DMEM with 10% FCS, and confluent cultures were routinely treated with TE buffer (0.05% trypsin, 0.02% EDTA). For force measurements, cells were treated with TE buffer (0.05% trypsin, 0.02% EDTA) and were seeded as single cells in a 60-mm ultra-low adhesion dishes (Corning) at 30,000 cells in 3 ml culture medium. After 18 hrs of suspension culture, cells were gently re-suspended in CO_2_-independent medium (Invitrogen) supplemented with 10% FCS and used immediately for SF measurement experiments.

### Measurement of separation force by pillar-pipette assay

A 60-mm ultra-low adhesion dish (Corning) was prepared by cutting a 2-cm window for pipette access. A PDMS block (Sylgard, Dow-Corning) with pillars 35 µm in diameter and 200 µm long arranged in a hexagonal pattern with a pitch of 400 µm was cut into a thin slice of 1-2 mm thickness and 1-cm long. The strip was further cut into 5 pieces of ∼2-mm thickness and stuck to the pre-prepared ultra-low adhesion dish in a staggered arrangement so that ∼20 micropillars at different x, y, z-levels were available to enable high-throughput. The dish was plasma treated to make the pillars hydrophilic (Harrick Plasma). During plasma treatment, a thin film of PDMS masked the area of the dish where the ultra-low property was to be maintained g. The pillars were coated with 50 µl fibronectin (Sigma-Aldrich) mixed with fluorescently labeled fibronectin (Cytoskeleton) in a ratio of 10:1 for 1 hr at 37 °C. The fibronectin was then washed with deionized water three times and the surface left to dry. The thin film of PDMS was removed before experiments.

Cells were manipulated at 37 °C with a micropipette that was held by a micromanipulator connected to a combined hydraulic/pneumatic system. Micropipettes were pulled (P-2000, Sutter Instrument) and cut with a microforge instrument (MF-900, Narishige) to an inner diameter of 4–6 µm. Cell doublets were visually observed under bright-field and epifluorescence to identify doublets with mature, bright junctions and a larger contact radius. One end of the doublet was attached to a PDMS pillar tip with the doublet axis parallel to the dish and perpendicular to the pillar’s axis (Fig. 2a,b and Supplementary Fig. 3d). After 5 min of attachment, a second pre-existing doublet was selected with the micropipette and brought in contact with the first doublet in a series arrangement (Fig. 2b). The new junction between the two doublets was established over 1 hr. The two doublets were separated from each other using a micropipette while the deflection in the pillar was imaged in bright-field (40× x 1.5, Nikon, Eclipse 2,000) at the rate of 1 image per sec. The SF required to separate the two doublets (after 1 hr contact) was quantified from the pillar deflection using the following equation (Supplementary Fig. 3d):

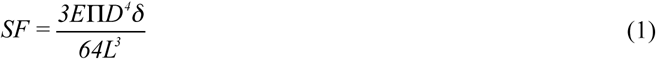

where *L* is the effective length of the pillar, *D* is the diameter, *E* is the Young’s Modulus of the PDMS material, and δ is the maximum pillar deflection before detaching the two doublets from each other (Supplementary Fig. 3d). Upon detaching the doublets, the two cells of the other doublet still attached to the pillar were separated using the micropipette, and the force required to separate the cells (pre-existing doublets) was determined. The results from 15 measurements from each group were used to obtain the mean SF for a specific contact time in at least three independent experiments.

### Dispase-based junctional protein tensor assay (D-JPT)

Channels made of UV curable polymer (NOA 73, Norland Products) were prepared using a PDMS stencil with the shape of the channel projecting out (100-500 µm high) placed upside-down on the glass-bottomed dish (IWAKI), and the low viscous UV curable polymer made to flow through the channels by capillarity effect and UV cured. The PDMS stencil was peeled off to expose the channel. The channel was washed three times by a jet of deionized water. Cells were seeded at 60% confluence in the rectangular glass-bottomed channel, 1-mm wide and 2.5-cm long with a central gradual constriction of 0.5 mm and 2.5 mm long (Fig. 5a). Cells were seeded within 3 h of making the channels to retain their hydrophobic characteristics to avoid cells attaching to the polymer. Upon reaching >95% confluence (Fig. 5b [fewer cells have been shown for ease of representation]), cells that had attached to the polymer, if any, were scrapped off with the back of a 10-µl pipette tip without damaging the cells inside the channel. Thereafter, a circular PDMS block with a central 0.5 cm × 0.5 cm window was inserted onto the dish tightly and the medium from within the window was pipetted out gently as to not damage the cell sheet in the channel (Fig. 5c). A 250-µl dispase solution (2.4 U/ml) was then gently pipetted into the opening and imaging at 20× was started in an inverted microscope (DMi8, Leica) acquiring 1 image per minute for 45 min to 1 hr.

Upon dispase treatment, the cells in the window region lifted off the substrate while constrained at the ends of the long-axis. As the edge of the cell sheet is not constrained in the short-axis, the sheet thinned out in the middle region due to the cells’ inherent cortical tension. The constriction in the middle of the channel makes the cells in that region more vulnerable to tension, causing the cells in this area to stretch at the expense of contraction of the cells on each side of the constriction (Fig. 5d).

### Cortical tension measurement and G-LISA assay

Cells were dissociated from confluent cultures by washing once with complete medium to remove loosely attached cells, and then gently flushing a spot of the monolayer to remove a patch of cells. This was done using a 1-ml pipette containing 1 ml CO_2_-independent medium supplemented with 10% FCS and flushing the spot three times with the same 1 ml of medium. The patch of removed cells was observed under the microscope to ensure that there were no cells in the area and thus the isolated cells were not a subpopulation of easily detachable cells with different rheology. The dissociated cells were gently pipetted 5 times to dissociate cell clumps before being deposited on a PLL-PEG-coated (SuSoS), glass-bottomed dish (IWAKI). A 1-cm window for pipette access was cut on the side of the glass-bottomed dish before cell seeding. Micropipettes were pulled as described earlier, and cut to a diameter ranging from 6 to 9 µm. Before each measurement, the pressure in the pipette was equilibrated with the pressure in the dish by monitoring the movement of cells when the pipette was brought closer to the cells. Cortical tension measurements were commenced 30 mins after the deposition of the cells onto the dish; this time delay allowed cells to become spherical at 37 °C. Single cells were gently aspirated into the pipette with a low pressure sufficient to grab the cell. The pressure was then gradually decreased until a threshold pressure was reached, at which point the cell formed a hemispherical protrusion into the pipette equal to the radius of the pipette (Fig. 3a). For measuring the cortical tension of doublets, the tip of the doublets was aspirated (Fig. 3b). The surface tension was computed using the Young-Laplace law *γ* = P_c_ / (2/R_p_ – 2/R_c_), where P_c_ is the negative pressure inside the pipette, R_c_ is the radius of the cell and R_p_ the radius of the pipette^30^. Experiments were performed within an environmental chamber maintained at 37 °C.

For G-LISA assay, to quantify active Rac1 and Cdc42 levels, cells were seeded at 50% confluence in a 10 cm cell culture dish. Upon reaching >95% confluence, scratch wounds were created using a 10-µl pipette tip, creating horizontal and vertical wounds at a spacing of ∼1 cm. The cultures were washed 3 times with 1x PBS (with calcium and magnesium) to remove the detached cells and incubated for a further 3 h to allow cells to start migrating, a process which is known to be regulated by Rho GTPases. The cells were then lysed, and G-LISA assay was performed as per the manufacturer’s instructions (Rac1 and Cdc42 G-LISA activation assay kits, Cytoskeleton).

### Preparation of wound-free gaps for cell migration assay

Wound gap (500 µm) masking inserts (Ibidi) were placed in the middle of the wells of 12-well cell culture-treated plates. Cells were seeded by pipetting 70 µl of cell suspension at 6 × 10^5^ cells/ml. After 18-24 h of culture, when the cells had reached >95% confluence, the inserts were carefully removed without disturbing the cells. The wells were gently washed three times with 1 ml PBS (with calcium and magnesium) to remove loose cells. Fresh medium (1 ml) was added to the culture, and the gap was imaged at different locations to acquire an image every 20 min using an inverted microscope (DMi8, Leica) equipped with an environmental chamber maintained at 37 °C and 5% CO_2_.

## Statistical analysis

All statistical analyses were performed with GraphPad Prism 5 software using an unpaired, two-tailed Student’s *t*-test. Data are presented as mean ± SD.

## Acknowledgements

We thank J. Ong, S. Nandi (IMCB) and T. Vicnesvari (SMART) for their excellent technical support with immunofluorescence staining, G-LISA assay and molecular cloning. We also thank K.H. Biswas (MBI) for sharing anti-vinculin antibody. This work was supported by MBI-Singapore seed funding (NRF grant). J.P.T acknowledges IMCB, A-STAR, core funding.

## Author contributions

D.M.K.A., RC.R., Y.S.C. and J.P.T. conceptualized the study. D.M.K.A.and R.C.R. prepared the plasmids and performed lentivirus-based transduction. D.M.K.A, V.V. and J.P.T. conceived the pillar-pipette assay. D.M.K.A. and J.P.T. conceived the D-JPT assay. D.M.K.A. performed the experiments and handled the data analysis. N.B. contributed the theoretical analysis. D.M.K.A. wrote the original draft. D.M.K.A., S.D., N.B., V.V. and J.P.T. wrote, reviewed and edited the manuscript. J.P.T. directed the research.

## Figure and table legends

**Supplementary Figure 1: Flow cytometry data** (BD Accuri C6) on cell groups previously FACS sorted based on overall GFP expression.

**Supplementary Figure 2: Confocal images showing cell surface cadherin-eGFP at the cell-substrate interface. (a)** E-cad-expressing cells. **(b)** 77EEE-expressing cells. **(c)** 777EE-expressing cells. **(d)** EE7EE-expressing cells. **(e)** 7777E-expressing cells. Scale bar, 15 µm.

**Supplementary Figure 3: Pillar-pipette assay. (a)** 3D Computer-Aided Design (CAD) model of 4 PDMS blocks glued to a 60-mm ultra-low dish, cut on both sides to facilitate pipette access. **(b)** Zoomed-in, top view of the dish with PDMS blocks showing pillars (∼200 µm long and ∼30 µm in diameter) parallel to the x-y plane of the dish and at different z-planes. Note the staggered arrangement of the blocks to facilitate a longer reach of the pipette entering from the left side; the pipette can reach the last pillar on the right extreme without disturbing other pillars along the way. **(c)** A closer look at individual pillars. Note the two pillars at different z-planes. **(d)** Bright-field images showing an undeflected, fibronectin-coated pillar (top), to the tip of which a 4-cell complex is attached and a micropipette accessing the free end of the cell complex. Note that the four cells, two pairs of preformed doublets after 1 hr contact are in series. The pillar deflects as the pipette pulls the cell to the left. The maximum pillar deflection (bottom) is used to calculate the SF using equation (1). Scale bar, 15 µm.

**Supplementary Figure 4: SF normalized against surface cadherin intensity. (a)** Histogram of SF at 1 hr contact, normalized against cell-surface cadherin intensity (E-cad, *n* = 15; 77EEE, *n* = 15; 777EE, *n* = 14; EE7EE, *n* = 15; 7777E, *n* = 15). **(b)** Histogram of SF required to separate pre-existing doublets (18 hrs), normalized against cell-surface cadherin intensity (E-cad, *n* = 15; 77EEE, *n* = 15; 777EE, *n* = 15; EE7EE, *n* = 14; 7777E, *n* = 15). Note that there is no significant difference between the raw and normalized SF values both for 1 hr and pre-existing (18 hrs) groups.

**Supplementary Figure 5: Flow cytometry, separation force (18 hrs) and wound-healing assay data of EEE7E and 77EE7 expressing S180 cells. (a)** Flow cytometry data showing GFP intensities of EEE7E, 77EE7 and S180(WT) cells. Note the low GFP intensity **(b)** SF of EEE7E (*n* = 7), 77EE7 (*n* = 6) and 7777E (*n* = 9) expressing cells. Note the SF of 77EE7 was not statistically different from 7777E cells. **(c)** Wound healing assay data showing the 77EE7 cells migrating faster than 777EE cells. The EEE7E cells migrated at a speed comparable to 7777E cells and not collectively.

**Supplementary Movie 1: S180 cells expressing strand-swap incompetent (Left) and X-dimer incompetent (Right) E-cad mutants**. Note that stable junctions were not formed in the S180 cells expressing the E-cad W2A mutant.

**Supplementary Movie 2: Pillar-pipette assay. (Left)** Representative movies from each group showing two pre-existing doublets adhering to each other at 1 hr contact and the 4-cell complex attached to a pillar tip. Note the deflection in pillars as the cells are pulled by the pipette. The SF values are listed on the left side. **(Right)** Representative movies from each group showing a pre-existing doublet attached to a pillar tip. The SF values are listed on the right side. The pillars are 36.7 µm in diameter and 188.7 µm long (full length not shown). The distance of cell attachment from the tip of the pillar is offset for calculating the effective length. Images were acquired at a rate of ∼1 frame per second.

**Supplementary Movie 3: Dispase-based junctional protein tensor assay. (Top)** E-cad-expressing S180 cells subjected to dispase treatment. Imaging commenced 5 min after the addition of dispase and was imaged for 1 hr, at a rate of 1 image per minute. Note that cells undergo significant strain while the cell-cell contacts are maintained. **(Middle)** 77EEE-expressing S180 cells subjected to similar treatment as in (e). Note that the cell sheet snaps immediately after the sheet is lifted and in lower strain levels as compared with E-cad-expressing cells in (a). **(Bottom)** 777EE-expressing S180 cells imaged for 15 min immediately after the addition of dispase. Although cell-cell junctions and a cell sheet are formed as for 77EEE-expressing cells, the 777EE-expressing cells are not able to maintain cell-cell contact in the absence of cell-substrate adhesion; they become single cells and cell clumps, unable to withstand the tension as a cell sheet. For all three frames, dotted circles show the area that is presented in higher magnification on the right. Scale bar, 100 µm.

**Supplementary Movie 4: D-JPT assay of S180 cells expressing EE7EE**. The previously cohesive cell sheet disintegrates into cell clumps, forming long membrane tethers in some cells. Note that the cortex of the cells forming tethers is rounded in the middle, indicative of higher cortical tension

**Supplementary Movie 5: Time-lapse confocal microscopy of S180 cells expressing cadherin-eGFP. (a)** E-cad-expressing cells **(b)** 77EEE-expressing cells **(c)** 777EE-expressing cells **(d)** EE7EE-expressing cells **(e)** 7777E-expressing cells, imaged for 6 min acquiring 1 image every 20 sec.

**Supplementary Movie 6: D-JPT assay of X-dimer incompetent E-cad K14E mutants**. Movie showing randomly oriented S180 cells expressing E-cad K14E mutants aligning along the axis of tension after dispase treatment. Refer to supplementary movie 2 for movie acquisition parameters.

**Supplementary Movie 7: Representative movies from each group showing 10 h of cell migration**. Images were acquired every 20 min until the wound-free gap disappears. Scale bar is 100 µm.

**Supplementary Movie 8:** Movies comparing cell migration in 77EE7 and EEE7E with E-cad, 77EEE and 7777E expressing S180 cells. Note the 77EE7 cells migrate collectively at a speed comparable to 77EEE cells. Also note the EEE7E cells migrating in a less cohesive manner and at speeds comparable to 7777E cells.

## References

1 Gumbiner, B. M. Cell adhesion: the molecular basis of tissue architecture and morphogenesis. Cell 84, 345–357 (1996).

2 Gumbiner, B. M. Regulation of cadherin-mediated adhesion in morphogenesis. Nat Rev Mol Cell Biol 6, 622–634, doi: 10.1038/nrm1699 (2005).

3 Takeichi, M. Morphogenetic roles of classic cadherins. Curr Opin Cell Biol 7, 619–627 (1995).

4 Hulpiau, P. & van Roy, F. New insights into the evolution of metazoan cadherins. Mol Biol Evol 28, 647–657, doi: 10.1093/molbev/msq233 (2011).

5 Boggon, T. J. et al. C-cadherin ectodomain structure and implications for cell adhesion mechanisms. Science 296, 1308–1313, doi: 10.1126/science.1071559 (2002).

6 Chu, Y. S. et al. Prototypical type I E-cadherin and type II cadherin-7 mediate very distinct adhesiveness through their extracellular domains. J Biol Chem 281, 2901–2910, doi: 10.1074/jbc.M506185200 (2006).

7 Halbleib, J. M. & Nelson, W. J. Cadherins in development: cell adhesion, sorting, and tissue morphogenesis. Genes Dev 20, 3199–3214, doi: 10.1101/gad.1486806 (2006).

8 Brochard-Wyart, F. & de Gennes, P. G. Unbinding of adhesive vesicles. C R Physique 4, 281–287, doi: 10.1016/S1631-0705(03)00048-3 (2003).

9 Patel, S. D. et al. Type II cadherin ectodomain structures: implications for classical cadherin specificity. Cell 124, 1255–1268, doi: 10.1016/j.cell.2005.12.046 (2006).

10 Harrison, O. J. et al. Two-step adhesive binding by classical cadherins. Nat Struct Mol Biol 17, 348–357, doi: 10.1038/nsmb.1784 (2010).

11 Sivasankar, S., Zhang, Y., Nelson, W. J. & Chu, S. Characterizing the initial encounter complex in cadherin adhesion. Structure 17, 1075–1081, doi: 10.1016/j.str.2009.06.012 (2009).

12 Ciatto, C. et al. T-cadherin structures reveal a novel adhesive binding mechanism. Nat Struct Mol Biol 17, 339–347, doi: 10.1038/nsmb.1781 (2010).

13 Rakshit, S., Zhang, Y., Manibog, K., Shafraz, O. & Sivasankar, S. Ideal, catch, and slip bonds in cadherin adhesion. Proc Natl Acad Sci U S A 109, 18815–18820, doi: 10.1073/pnas.1208349109 (2012).

14 Leckband, D. & Sivasankar, S. Cadherin recognition and adhesion. Curr Opin Cell Biol 24, 620–627, doi: 10.1016/j.ceb.2012.05.014 (2012).

15 Thiery, J. P., Engl, W., Viasnoff, V. & Dufour, S. Biochemical and biophysical origins of cadherin selectivity and adhesion strength. Curr Opin Cell Biol 24, 614–619, doi: 10.1016/j.ceb.2012.06.007 (2012).

16 Sivasankar, S., Brieher, W., Lavrik, N., Gumbiner, B. & Leckband, D. Direct molecular force measurements of multiple adhesive interactions between cadherin ectodomains. Proc Natl Acad Sci U S A 96, 11820–11824 (1999).

17 Sivasankar, S., Gumbiner, B. & Leckband, D. Direct measurements of multiple adhesive alignments and unbinding trajectories between cadherin extracellular domains. Biophys J 80, 1758–1768, doi: 10.1016/S0006-3495(01)76146-2 (2001).

18 Zhu, B. et al. Functional analysis of the structural basis of homophilic cadherin adhesion. Biophys J 84, 4033–4042, doi: 10.1016/S0006-3495(03)75129-7 (2003).

19 Perez, T. D. & Nelson, W. J. Cadherin adhesion: mechanisms and molecular interactions. Handb Exp Pharmacol, 3–21, doi: 10.1007/978-3-540-68170-0_1 (2004).

20 Fichtner, D. et al. Covalent and density-controlled surface immobilization of E-cadherin for adhesion force spectroscopy. PLoS One 9, e93123, doi: 10.1371/journal.pone.0093123 (2014).

21 Bayas, M. V., Leung, A., Evans, E. & Leckband, D. Lifetime measurements reveal kinetic differences between homophilic cadherin bonds. Biophys J 90, 1385–1395, doi: 10.1529/biophysj.105.069583 (2006).

22 Chappuis-Flament, S., Wong, E., Hicks, L. D., Kay, C. M. & Gumbiner, B. M. Multiple cadherin extracellular repeats mediate homophilic binding and adhesion. J Cell Biol 154, 231–243 (2001).

23 Chien, Y. H. et al. Two stage cadherin kinetics require multiple extracellular domains but not the cytoplasmic region. J Biol Chem 283, 1848–1856, doi: 10.1074/jbc.M708044200 (2008).

24 Shi, Q., Maruthamuthu, V., Li, F. & Leckband, D. Allosteric cross talk between cadherin extracellular domains. Biophys J 99, 95–104, doi: 10.1016/j.bpj.2010.03.062 (2010).

25 Liu, R., Wu, F. & Thiery, J. P. Remarkable disparity in mechanical response among the extracellular domains of type I and II cadherins. J Biomol Struct Dyn 31, 1137–1149, doi: 10.1080/07391102.2012.726530 (2013).

26 Chu, Y. S. et al. Force measurements in E-cadherin-mediated cell doublets reveal rapid adhesion strengthened by actin cytoskeleton remodeling through Rac and Cdc42. J Cell Biol 167, 1183–1194, doi: 10.1083/jcb.200403043 (2004).

27 Maitre, J. L. et al. Adhesion functions in cell sorting by mechanically coupling the cortices of adhering cells. Science 338, 253–256, doi: 10.1126/science.1225399 (2012).

28 Borghi, N. & James Nelson, W. Intercellular adhesion in morphogenesis: molecular and biophysical considerations. Curr Top Dev Biol 89, 1–32, doi: 10.1016/S0070-2153(09)89001-7 (2009).

29 Winklbauer, R. Cell adhesion strength from cortical tension - an integration of concepts. J Cell Sci 128, 3687–3693, doi: 10.1242/jcs.174623 (2015).

30 Maitre, J. L., Niwayama, R., Turlier, H., Nedelec, F. & Hiiragi, T. Pulsatile cell-autonomous contractility drives compaction in the mouse embryo. Nat Cell Biol 17, 849–855, doi: 10.1038/ncb3185 (2015).

31 Youssef, J., Nurse, A. K., Freund, L. B. & Morgan, J. R. Quantification of the forces driving self-assembly of three-dimensional microtissues. Proc Natl Acad Sci U S A 108, 6993–6998, doi: 10.1073/pnas.1102559108 (2011).

32 Stirbat, T. V. et al. Fine tuning of tissues’ viscosity and surface tension through contractility suggests a new role for alpha-catenin. PLoS One 8, e52554, doi: 10.1371/journal.pone.0052554 (2013).

33 Engl, W., Arasi, B., Yap, L. L., Thiery, J. P. & Viasnoff, V. Actin dynamics modulate mechanosensitive immobilization of E-cadherin at adherens junctions. Nat Cell Biol 16, 587–594, doi: 10.1038/ncb2973 (2014).

34 Arnold, K., Bordoli, L., Kopp, J. & Schwede, T. The SWISS-MODEL workspace: a web-based environment for protein structure homology modelling. Bioinformatics 22, 195–201, doi: 10.1093/bioinformatics/bti770 (2006).

35 Tabdili, H. et al. Cadherin point mutations alter cell sorting and modulate GTPase signaling. J Cell Sci 125, 3299–3309, doi: 10.1242/jcs.087395 (2012).

36 Bi, D., Lopez, J. H., Schwarz, J. M. & Manning, M. L. A density-independent rigidity transition in biological tissues. Nature Physics 11, 1074–1079, doi: DOI: 10.1038/NPHYS3471 (2015).

37 Thomas, W. A. et al. alpha-Catenin and vinculin cooperate to promote high E-cadherin-based adhesion strength. J Biol Chem 288, 4957–4969, doi: 10.1074/jbc.M112.403774 (2013).

38 Yonemura, S., Wada, Y., Watanabe, T., Nagafuchi, A. & Shibata, M. alpha-Catenin as a tension transducer that induces adherens junction development. Nat Cell Biol 12, 533–542, doi: 10.1038/ncb2055 (2010).

39 Liwosz, A., Lei, T. & Kukuruzinska, M. A. N-glycosylation affects the molecular organization and stability of E-cadherin junctions. J Biol Chem 281, 23138–23149, doi: 10.1074/jbc.M512621200 (2006).

40 Vagin, O., Tokhtaeva, E., Yakubov, I., Shevchenko, E. & Sachs, G. Inverse correlation between the extent of N-glycan branching and intercellular adhesion in epithelia. Contribution of the Na,K-ATPase beta1 subunit. J Biol Chem 283, 2192–2202, doi: 10.1074/jbc.M704713200 (2008).

41 Brasch, J. et al. Structure and binding mechanism of vascular endothelial cadherin: a divergent classical cadherin. J Mol Biol 408, 57–73, doi: 10.1016/j.jmb.2011.01.031 (2011).

42 le Duc, Q. et al. Vinculin potentiates E-cadherin mechanosensing and is recruited to actin-anchored sites within adherens junctions in a myosin II-dependent manner. J Cell Biol 189, 1107–1115, doi: 10.1083/jcb.201001149 (2010).

43 Martinez-Rico, C., Pincet, F., Thiery, J. P. & Dufour, S. Integrins stimulate E-cadherin-mediated intercellular adhesion by regulating Src-kinase activation and actomyosin contractility. J Cell Sci 123, 712–722, doi: 10.1242/jcs.047878 (2010).

44 Guo, H. B., Johnson, H., Randolph, M. & Pierce, M. Regulation of homotypic cell-cell adhesion by branched N-glycosylation of N-cadherin extracellular EC2 and EC3 domains. J Biol Chem 284, 34986–34997, doi: 10.1074/jbc.M109.060806 (2009).

45 Langer, M. D., Guo, H., Shashikanth, N., Pierce, J. M. & Leckband, D. E. N-glycosylation alters cadherin-mediated intercellular binding kinetics. J Cell Sci 125, 2478–2485, doi: 10.1242/jcs.101147 (2012).

46 Decave, E., Garrivier, D., Brechet, Y., Bruckert, F. & Fourcade, B. Peeling process in living cell movement under shear flow. Phys Rev Lett 89, 108101, doi: 10.1103/PhysRevLett.89.108101 (2002).

47 Harrison, O. J. et al. The extracellular architecture of adherens junctions revealed by crystal structures of type I cadherins. Structure 19, 244–256, doi: 10.1016/j.str.2010.11.016 (2011).

48 Borghi, N. & Brochard-Wyart, F. Tether extrusion from red blood cells: integral proteins unbinding from cytoskeleton. Biophys J 93, 1369–1379, doi: 10.1529/biophysj.106.087908 (2007).

49 Tabdanov, E., Borghi, N., Brochard-Wyart, F., Dufour, S. & Thiery, J. P. Role of E-cadherin in membrane-cortex interaction probed by nanotube extrusion. Biophys J 96, 2457–2465, doi: 10.1016/j.bpj.2008.11.059 (2009).

50 Yamada, S. & Nelson, W. J. Localized zones of Rho and Rac activities drive initiation and expansion of epithelial cell-cell adhesion. J Cell Biol 178, 517–527, doi: 10.1083/jcb.200701058 (2007).

51 Hidalgo-Carcedo, C. et al. Collective cell migration requires suppression of actomyosin at cell-cell contacts mediated by DDR1 and the cell polarity regulators Par3 and Par6. Nat Cell Biol 13, 49–58, doi: 10.1038/ncb2133 (2011).

52 Toret, C. P., Collins, C. & Nelson, W. J. An Elmo-Dock complex locally controls Rho GTPases and actin remodeling during cadherin-mediated adhesion. J Cell Biol 207, 577–587, doi: 10.1083/jcb.201406135 (2014).

53 Jasaitis, A., Estevez, M., Heysch, J., Ladoux, B. & Dufour, S. E-cadherin-dependent stimulation of traction force at focal adhesions via the Src and PI3K signaling pathways. Biophys J 103, 175–184, doi: 10.1016/j.bpj.2012.06.009 (2012).

54 Bialucha, C. U. et al. Discovery and Optimization of HKT288, a Cadherin-6-Targeting ADC for the Treatment of Ovarian and Renal Cancers. Cancer Discov 7, 1030–1045, doi: 10.1158/2159-8290.CD-16-1414 (2017).

55 Emsley, P., Lohkamp, B., Scott, W. G. & Cowtan, K. Features and development of Coot. Acta Crystallogr D Biol Crystallogr 66, 486–501, doi: 10.1107/S0907444910007493 (2010).

