## supplementary figures for "Extracellular domains of E-cadherin determine key mechanical phenotypes of an epithelium through cell- and non-cell-autonomous outside-in signalling"

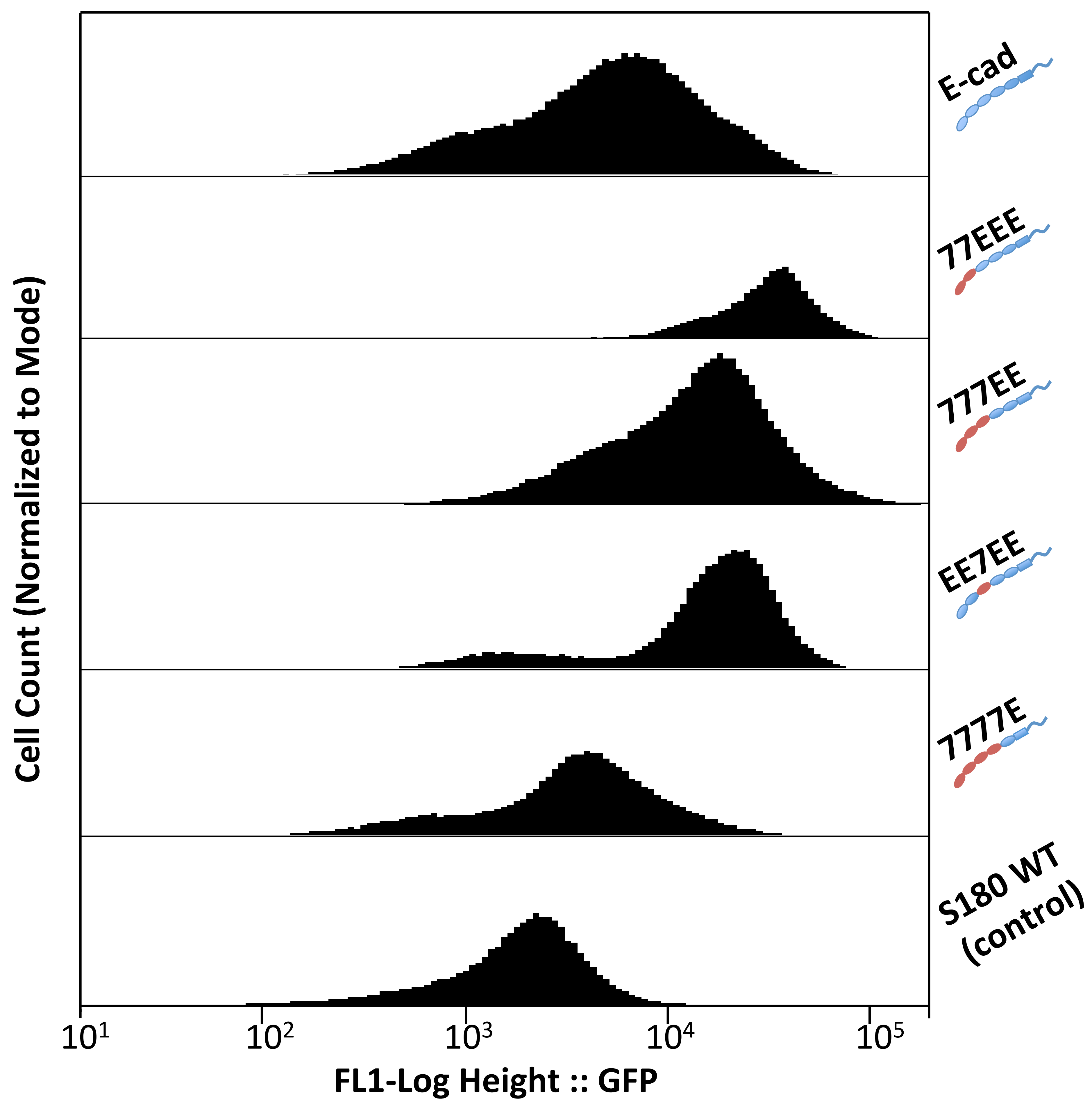


**Supplementary Figure 1: Flow cytometry data.**

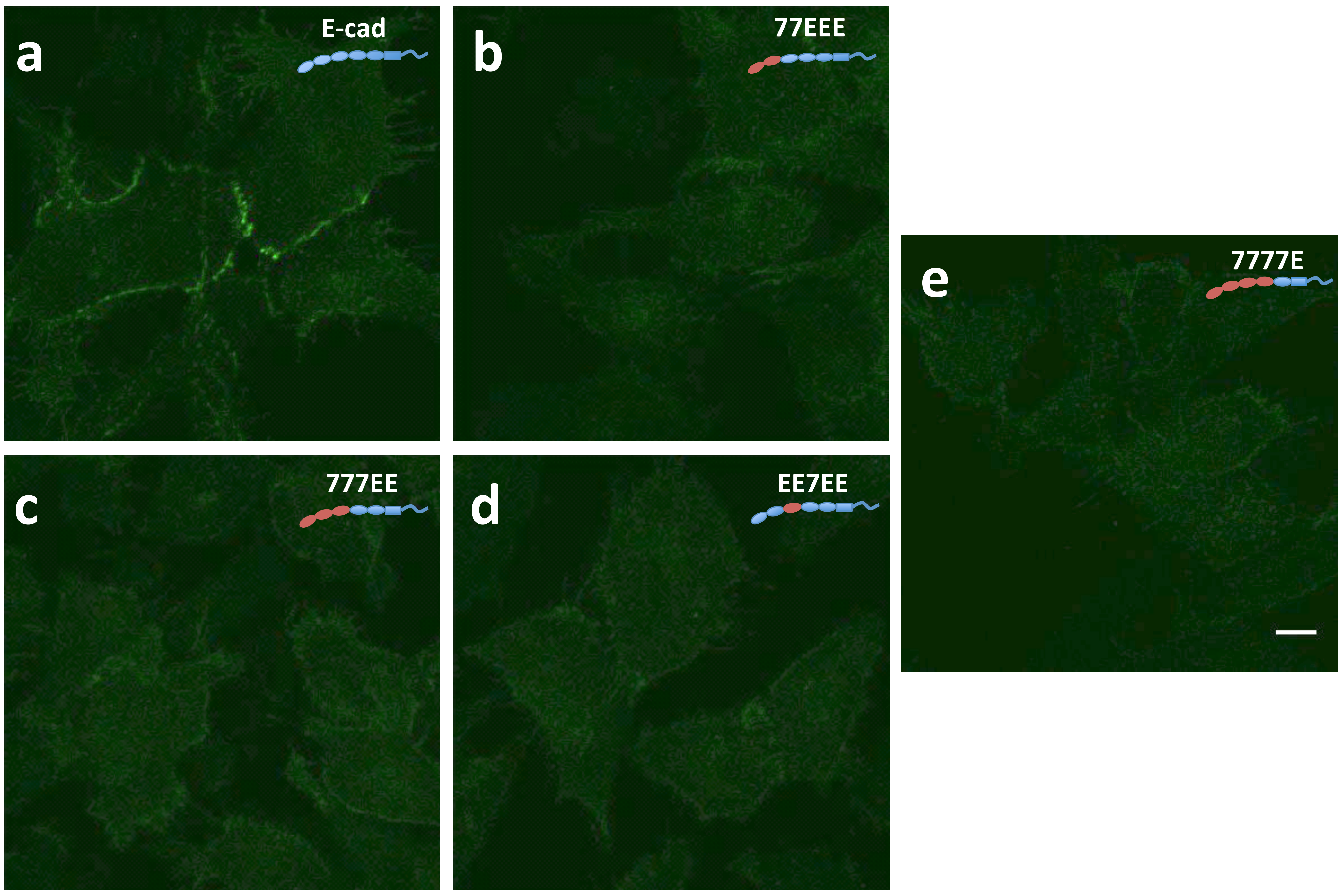


**Supplementary Figure 2: Confocal images showing cell surface cadherin-eGFP at the cell-substrate interface.**

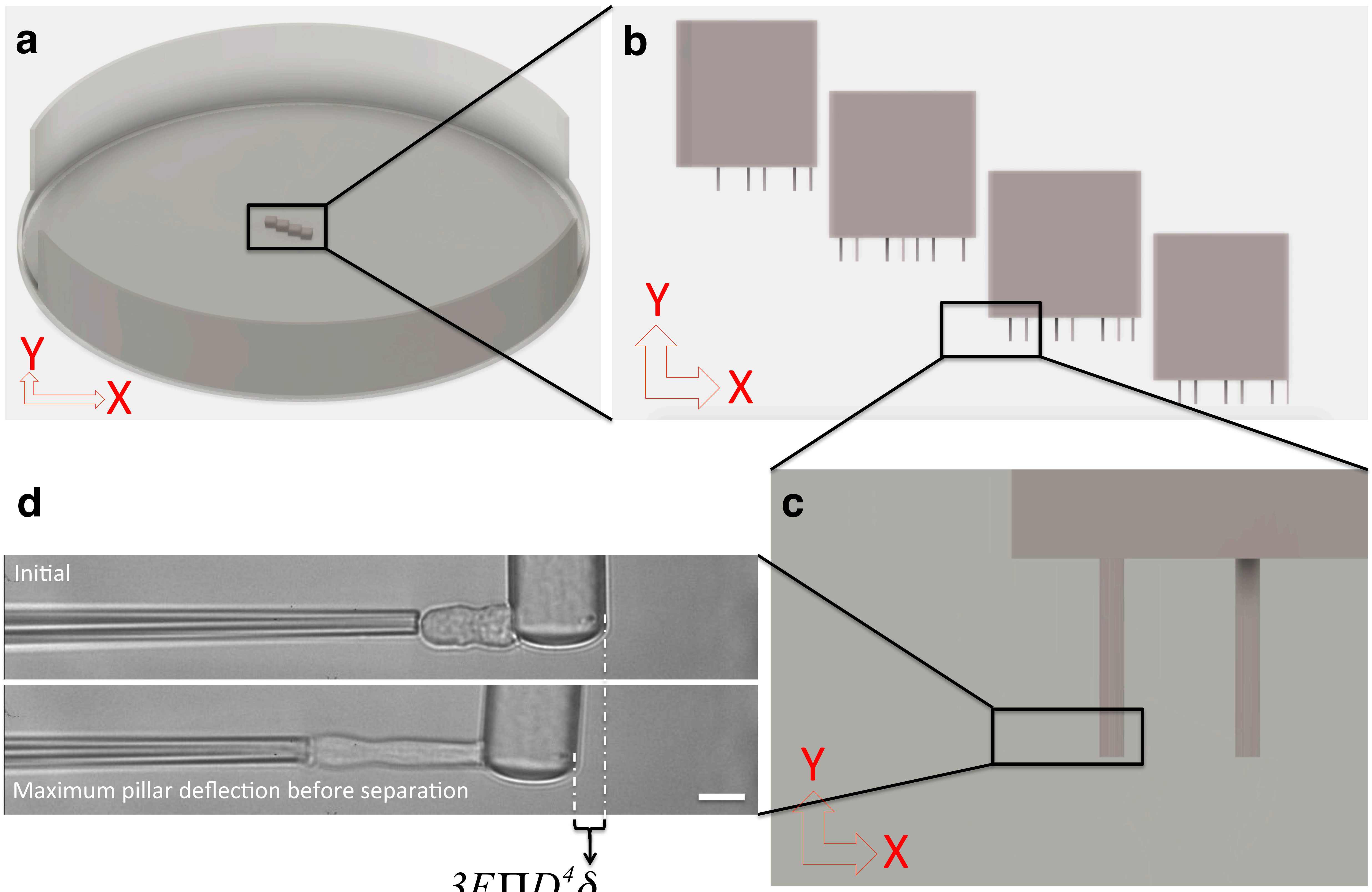


$$Separation\ Force = \frac{3E\Pi D^4\delta}{64L^3} \quad \text{— Eq. (1)}$$

**Supplementary Figure 3: Pillar-pipette assay.**

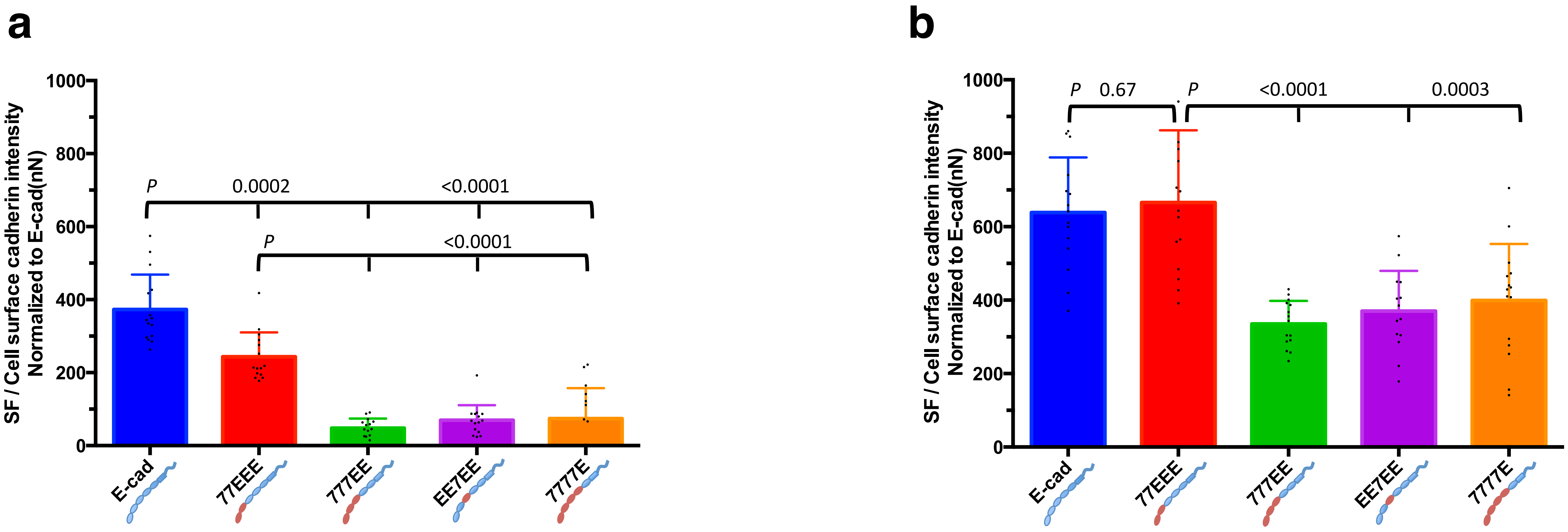


**Supplementary Figure 4: SF normalized against surface cadherin intensity.**

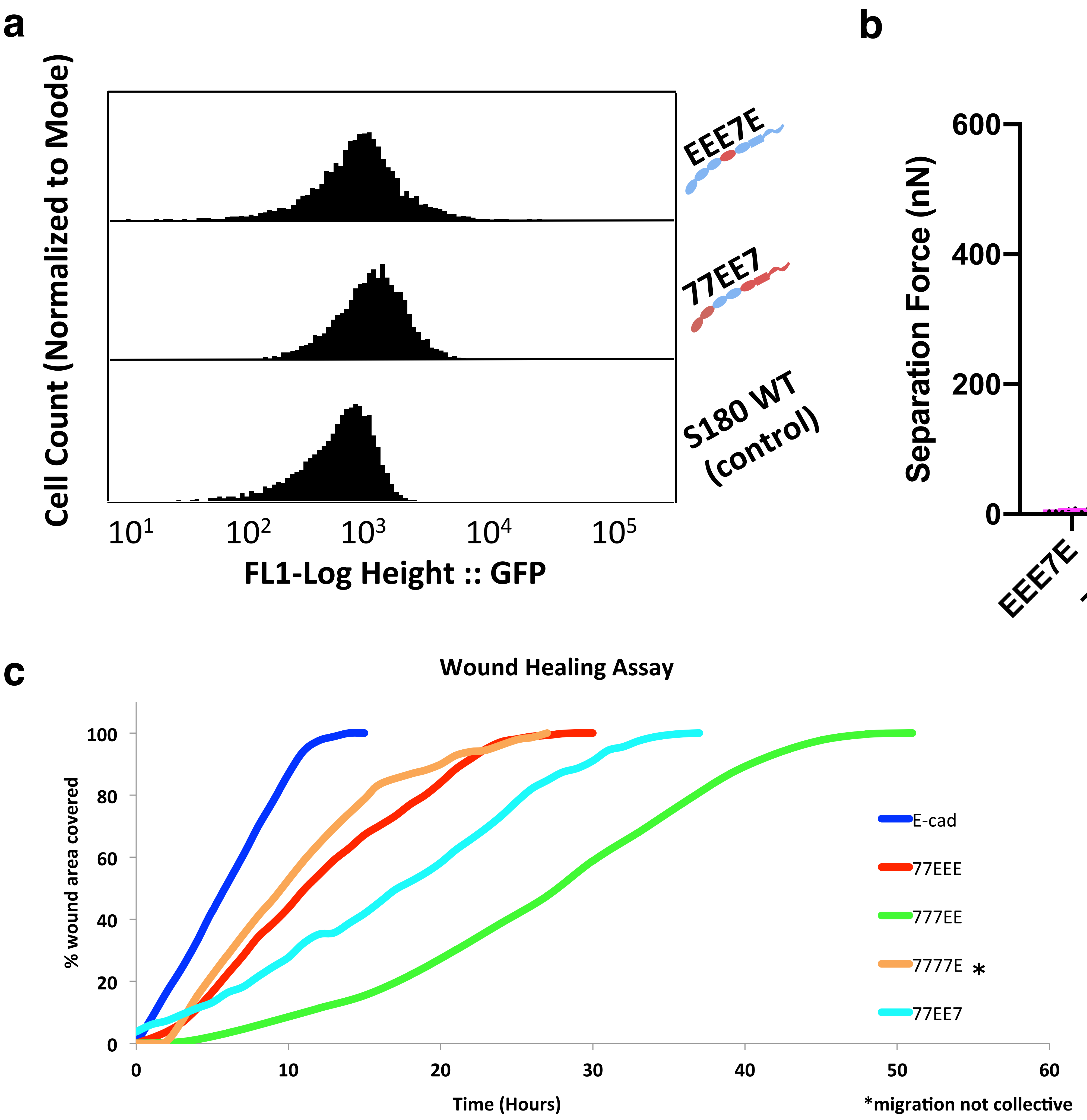


**Supplementary Figure 5: Flow cytometry data of EEE7E and 77EE7, separation force (18 hrs) and wound healing assay data of EEE7E and 77EE7 expressing S180 cells.**
